# Transcriptional regulation of neural stem cell expansion in adult hippocampus

**DOI:** 10.1101/2021.07.14.452351

**Authors:** Nannan Guo, Kelsey D. McDermott, Yu-Tzu Shih, Haley Zanga, Debolina Ghosh, Charlotte Herber, James Coleman, Alexia Zagouras, William R. Meara, Lai Ping Wong, Ruslan Sadreyev, J. Tiago Gonçalves, Amar Sahay

**Author notes:** Corresponding author: Amar Sahay. These authors contributed equally (Kelsey D. McDermott, Yu-Tzu Shih).

## Abstract

Experience governs neurogenesis from radial-glial neural stem cells (RGLs) in the adult hippocampus to support memory. Transcription factors in RGLs integrate physiological signals to dictate self-renewal division mode. Whereas asymmetric RGL divisions drive neurogenesis during favorable conditions, symmetric divisions prevent premature neurogenesis while amplifying RGLs to anticipate future neurogenic demands. The identities of transcription factors regulating RGL symmetric self-renewal, unlike those that regulate RGL asymmetric self-renewal, are not known. Here, we show that the transcription factor Kruppel-like factor 9 (*Klf9*) is elevated in quiescent RGLs and inducible, deletion of *Klf9* promotes RGL activation state. Clonal analysis and longitudinal intravital 2-photon imaging directly demonstrate that Klf9 functions as a brake on RGL symmetric self-renewal. *In vivo* translational profiling of RGLs lacking Klf9 generated a blueprint of RGL symmetric self-renewal for stem cell community. Together, these observations identify Klf9 as a transcriptional regulator of neural stem cell expansion in the adult hippocampus.

## Introduction

In the adult mammalian brain, radial glial neural stem cells (RGLs) in the dentate gyrus subregion of the hippocampus give rise to dentate granule cells and astrocytes (*1-9*), a process referred to as adult hippocampal neurogenesis (*10-18*). Adult-born dentate granule cells integrate into hippocampal circuitry by remodeling the network and ultimately contribute to hippocampal dependent learning and memory and regulation of emotion (*7, 19, 20*). Levels of adult hippocampal neurogenesis are highly sensitive to experience (*21, 22*) suggesting that neurogenesis may represent an adaptive mechanism by which hippocampal dependent memory functions are optimized in response to environmental demands. Essential to this adaptive flexibility is the capacity of RGLs to balance long-term maintenance with current or future demands for neurogenesis (“anticipatory neurogenesis”) in response to distinct physiological signals (*5, 21-24*).

Depending on environmental conditions, RGLs make decisions to stay quiescent or self-renew asymmetrically or symmetrically. Whereas enriching experiences (eg: complex environments, exploration, socialization) bias RGLs towards asymmetric divisions to generate astrocytes and neurons (*23, 25*), unfavorable conditions promote RGL quiescence (eg: chronic stress, aging) or symmetric self-renewal to support neural stem cell expansion at the expense of neurogenesis (eg: social isolation, seizures, aging) (*23, 26, 27*). Asymmetric self-renewal is thought to be the predominant mode of RGL division in the adult hippocampus and it ensures maintenance of RGL numbers while supporting current neurogenic demands (*9, 22*). Conversely, symmetric self-renewal decouples RGL divisions from differentiation and is thought to serve distinct functions. First, symmetric divisions prevent premature differentiation of RGLs in a non-permissive or unhealthy niche, and consequently, avert aberrant integration of adult-born dentate granule cells detrimental to hippocampal functions (*27, 28*). As such, RGL amplification anticipates future demands for neurogenesis upon return to favorable conditions. Second, RGL expansion may represent an efficient mechanism to replenish the adult RGL pool after injury. Third, symmetric stem cell divisions maybe more efficient than asymmetric divisions for long-term maintenance since fewer divisions are required to maintain RGL numbers. Furthermore, symmetric divisions may be associated with a lower rate of mutations and reduced replicative aging (*29*).

Extracellular physiological signals recruit transcription factors (TFs) within adult hippocampal RGLs to execute quiescence-activation decisions and symmetric or asymmetric self-renewal divisions (*22, 30, 31*). A growing number of transcriptional regulators of quiescence and asymmetric (neurogenic or astrogenic) stem cell renewal has been identified (*32-36*). Deletion of such factors results in loss of RGL quiescence, increased neurogenesis and ultimately, differentiation-coupled depletion of the RGL pool. In sharp contrast, the identities of TFs that regulate RGL expansion have remained elusive. Here, we report that expression of the ubiquitously expressed TF, Kruppel-like factor 9 (*Klf9)*, a regulator of dendritic and axonal plasticity in post-mitotic neurons (*37, 38*), is elevated in non-dividing RGLs compared to dividing RGLs. Inducible genetic upregulation of *Klf9* in RGLs and progenitors decreased activation, whereas conditional cell-autonomous deletion of *Klf9* in RGLs promoted an activated state. Clonal lineage tracing and longitudinal two-photon imaging of adult hippocampal RGLs *in vivo* directly demonstrated a role for *Klf9* as a brake on symmetric self-renewal. *In vivo* translational profiling of RGLs generated a molecular blueprint for RGL expansion in the adult hippocampus: we found that loss of Klf9 in RGLs results in downregulation of a program of quiescence associated factors and upregulation of genetic (mitogen, notch) and metabolic (fatty acid oxidation and lipid signaling) programs underlying RGL symmetric self-renewal. Together, these data identify Klf9 as a transcriptional regulator of neural stem cell expansion in the adult hippocampus. Our study contributes to an emerging framework for how experiential signals may toggle a balance of transcriptional regulators of symmetric and asymmetric self-renewal of RGLs to amplify neural stem cells or asymmetrically divide and generate neurons and astrocytes.

## Results

### Inducible *Klf9* loss promotes RGL activation

To characterize *Klf9* expression in RGLs in the adult dentate gyrus, we bred *Klf9*-*LacZ* knock-in reporter mice (*39*) with a Nestin GFP transgenic mouse line in which Nestin+RGLs are genetically labeled with GFP (*40*). Quantification of *Klf9* expression based on LacZ intensities in *Klf9* ^*LacZ/+*^ mice revealed enrichment in quiescent RGLs relative to activated RGLs (MCM2+)(One-way ANOVA, F=17.07, p=0.003)(**Figures 1A-B**). MCM2 expression captures activated cells that have exited quiescence. To refine this estimation that is based on a surrogate (LacZ) of *Klf9* expression within the RGL compartment, we performed Fluorescence in situ hybridization (FISH) using a *Klf9* specific riboprobe and immunohistochemistry for GFP and BrdU on adult hippocampal sections obtained from *Klf9* ^*+/+* or *LacZ/LacZ*^; Nestin GFP transgenic mice perfused 2 hours following a BrdU pulse (One-way ANOVA, F=5.6, p=0.04)(**Figures 1C-E**). No signal was detected with FISH using the *Klf9* riboprobe on brain sections from Klf9 ^*LacZ/LacZ*^ mice thus conveying specificity of the riboprobe (**Figure 1D**). Quantification of *Klf9* transcripts using Image J revealed significantly enriched expression in quiescent vs. activated (Brdu+Nestin GFP+) RGLs (**Figures 1C and 1E**).

**Figure 1.**
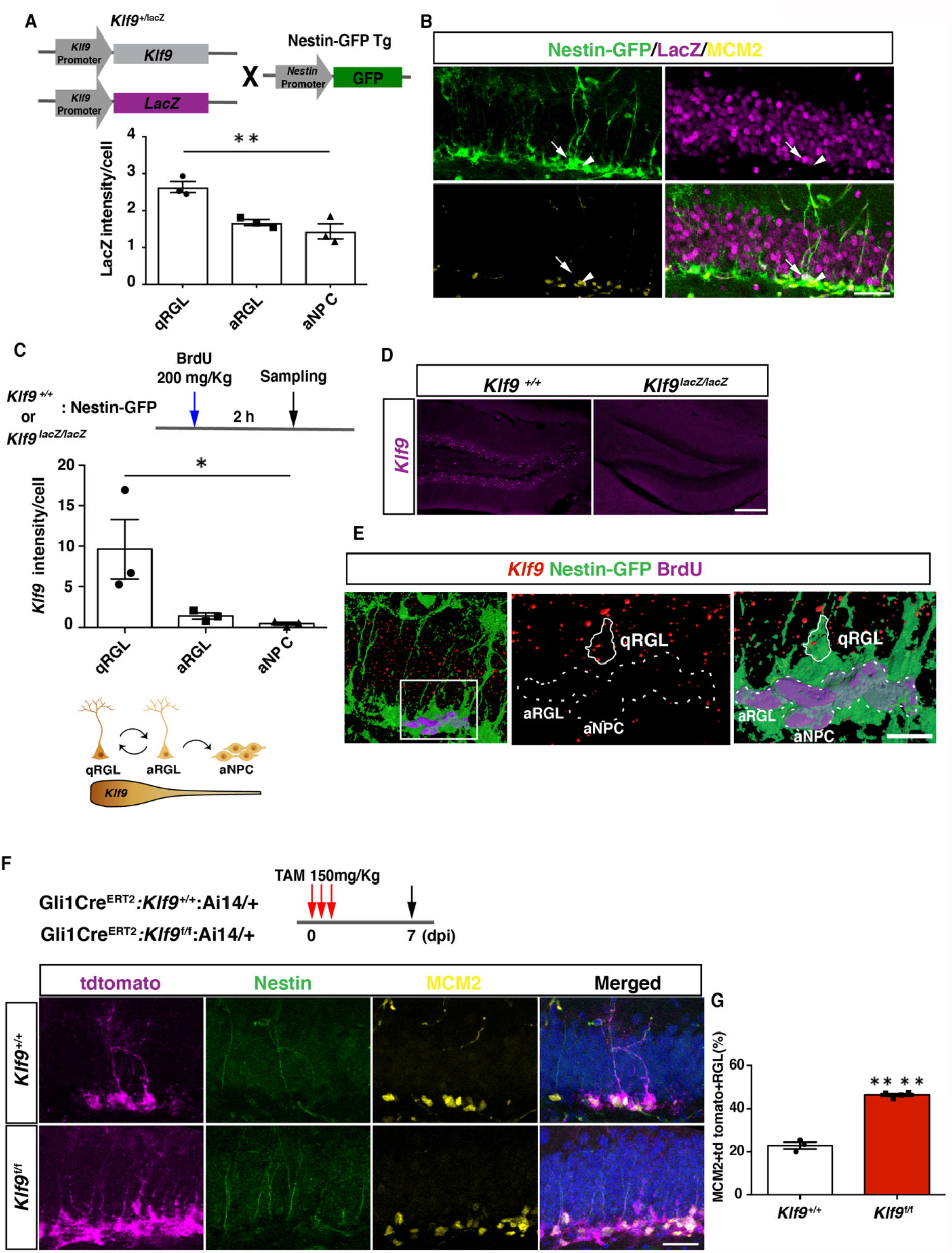
*Klf9* is elevated in non-dividing RGLs and loss of Klf9 promotes RGL activation. (A-B) *Klf9* expression inferred from LacZ expression intensity in quiescent RGLs, qRGL (GFP+MCM2-with radial process, arrows), activated RGL, aRGL (GFP+MCM2+ with radial process, arrowheads) and activated neural progenitors, aNPCs (GFP+MCM2+ without a radial process) in *Klf9* ^*LacZ/+* or *LacZ/LacZ*^; Nestin GFP transgenic. qRGLs exhibit higher *Klf9* expression than aRGLs and aNPCs. n=3 mice/group. (C-E) Fluorescence *in situ* hybridization using a *Klf9* specific riboprobe and immunohistochemistry for GFP and BrdU on adult hippocampal sections obtained from *Klf9* ^*LacZ/+* or *LacZ/LacZ*^; Nestin GFP transgenic mice. (D) Specificity of riboprobe established by detection of *Klf9* expression in dentate gyrus of *Klf9* ^*+/+*^ but not in *Klf9* ^*LacZ/LacZ*^ mice. (C, E) *Klf9* is expressed in qRGLs but not in dividing (BrdU+) RGLs or aNPCs. n=3 mice/group. (F-G) Inducible deletion of *Klf9* in Gli1+ RGLs in adult mice (Gli1 Cre^ERT2^*:Klf9*^+/+^:Ai14 vs. Gli1 Cre^ERT2^*:Klf9*^f/f^:Ai14) results in increased RGL activation (percentage of MCM2+tdTomato+Nestin+RGLs). n=3, 4 mice/group. Data are represented as mean ± SEM. * p<0.05, ** p<0.01, **** p<0.0001. Scale Bar Figures 1B, 1F: 50 μm, Figure 1D 250 μm, Figure 1E 20 μm. See also corresponding Supplementary Figures 1 and 2.

We next asked what happens when we delete *Klf9* in adult hippocampal RGLs. To address this question, we engineered *Klf9* conditional mutant mice (*Klf9*^*f/f*^) to cell-autonomously delete *Klf9* in RGLs. We first validated our *Klf9*^*f/f*^ mouse line by crossing it with the POMC-Cre mouse line that drives recombination in the dentate gyrus. ISH on hippocampal sections from POMC-Cre: *Klf9*^f/f^ revealed salt and pepper expression of *Klf9* in the dentate gyrus consistent with the established pattern of POMC-Cre dependent recombination in the dentate gyrus (*41*). No signal was detected by in situ hybridization using the *Klf9* riboprobe on brain sections from *Klf9* ^*LacZ/LacZ*^ mice thus conveying specificity of the riboprobe (**Figures S1A-C**). We bred *Klf9*^*f/f*^ mice with (GLI-Kruppel family member 1) Gli1 Cre^ERT2^ to recombine *Klf9* (deletion of exon1) in hippocampal RGLs (**Figures 1F-G**). We chose the Gli1 Cre^ERT2^ driver line because population-based lineage tracing and chronic *in vivo* imaging suggests that Gli1Cre^ERT2^ labeled RGLs contribute to long-term maintenance and self-renewal (*3, 42*). We next generated *Klf9*^f/f or +/+^ mice harboring a Gli1 Cre^ERT2^ allele and a Cre-reporter allele (Ai14, B6;129S6-*Gt(ROSA)26Sor*^*tm14(CAG-tdTomato)Hze*^/J)(*43*) to indelibly label Gli1-positive RGLs and their progeny (**Figure 1F)**. We induced *Klf9* recombination and tdTomato expression in RGLs of adult (2 months old) Gli1 Cre^ERT2^; *Klf9*^f/f or +/+^; Ai14 mice and processed brain sections for Nestin, tdTomato, and MCM2 immunohistochemistry 7 days post injection (7dpi) to quantify activated RGLs (**Figures 1F-G**). We found that conditional deletion of *Klf9* in Gli1 Cre^ERT2^ targeted adult hippocampal RGLs significantly increased the fraction of activated RGLs (% of MCM2+tdTomato+RGLs)(**Figure 1G**)(Unpaired t-tests, Figure 1G p=0.02, Figure 1I p<0.0001). Complementing these results, we demonstrated that genetic overexpression of *Klf9* in activated hippocampal neural stem cells and progenitors of adult Sox1 tTA; teto Klf9 mice (*38, 44*) significantly decreased the fraction of activated and dividing cells (**Figures S2A-D**). Together, these data demonstrate that *Klf9* expression is enriched in quiescent RGLs and that loss of *Klf9* expression in RGLs either promotes or maintains an activated state of RGLs in the adult hippocampus.

### *Klf9* deletion in RGLs produces supernumerary RGL clones

We next asked how Klf9 loss of function in RGLs affects self-renewal division mode. Population level lineage tracing experiments at short-term chase time points suggested that Klf9 loss in Gli1+ RGLs increased RGL numbers (data not shown). However, analysis of neural stem cell dynamics at the population level is encumbered by changes in numbers of labeled progeny overtime (*5, 42*). The challenges of interpreting population-level analysis are exacerbated because *Klf9* is also expressed in immature adult-born neurons and mature dentate granule cells. As such, changes in numbers of labeled descendants following loss of Klf9 in RGLs makes population-level lineage tracing difficult to interpret. Therefore, to directly investigate whether loss of *Klf9* in RGLs results in neural stem cell expansion at a single clone level, we performed *in vivo* clonal analysis in adult Gli1 Cre^ERT2^; *Klf9*^f/f or +/+^; Ai14 mice shortly after low dose Tamoxifen adminsitration. Single dose of TAM at 50mg/Kg body weight permitted sparse labeling of single tdTomato+RGLs and visualization of labeled single RGL clones and their individual constituents. Analysis of clonal composition at 7dpi revealed a significantly greater fraction of multi-RGL containing clones and a smaller fraction of single RGL containing clones 7 day post *Klf9* deletion in RGLs (**Figures 2A-D**)(Figure 2B Two-way ANOVA, Genotype X Cell-type p<0.0001, Bonferroni *post hoc Klf9* ^+/+^ vs. *Klf9*^f/f.^ 1RGL n.s., 2+RGLs p<0.0001, 1RGLs+ p<0.0001, No RGL n.s. Figure 2D, left panel, Two-way ANOVA, Genotype X Cell-type p<0.0001, Bonferroni *post hoc Klf9* ^+/+^ vs. *Klf9*^f/f.^ 2+RGLs p=0.09, 2RGLs+P+A p<0.0001, 2RGLs+P n.s. Figure 2D, right panel, Two-way ANOVA, Genotype X Cell-type p=0.09, Bonferroni *post hoc Klf9* ^+/+^ vs. *Klf9*^f/f.^ 1RGL n.s., 1RGL+P+A p=0.01, 1RGLs+P p=0.01). Many of the multi-RGL containing clones also comprised of neural progenitors and astrocytes suggesting that loss of *Klf9* biases RGL expansion but does not prevent RGL differentiation into progeny (**Figure 2D**). To corroborate these findings and address any potential confound introduced by bias in the Ai14 genetic lineage tracer, we performed clonal analysis at 7dpi using a different lineage tracing reporter, mTmG (*t(ROSA)26Sor*^*tm4(ACTB-tdTomato,-EGFP)Luo*^/J)(*45*). Our analysis demonstrated a significant increase in multi-RGL clones and decrease in single RGL clones derived from RGLs lacking *Klf9* in Gli1 Cre^ERT2^; *Klf9* ^f/f^mTmG mice (**Figures 2E-F**)(Figure 2F, Two-way ANOVA, Genotype X Cell-type p<0.0001, Bonferroni *post hoc Klf9* ^+/+^ vs. *Klf9*^f/f.^ 1RGL n.s., 2+RGLs p=0.0001, 1+RGLs p=0.03, No RGLs n.s). The lack of a difference in single RGL clones suggests that loss of Klf9 maintains the activated state associated with increased symmetric divisions. Alternatively, it may reflect a floor effect in the assay that occludes detection of a decrease in number. These findings provide evidence for *Klf9* in cell-autonomous regulation of RGL expansion and are *suggestive* of a role for Klf9 in inhibition of symmetric self-renewal of RGLs.

**Figure 2.**
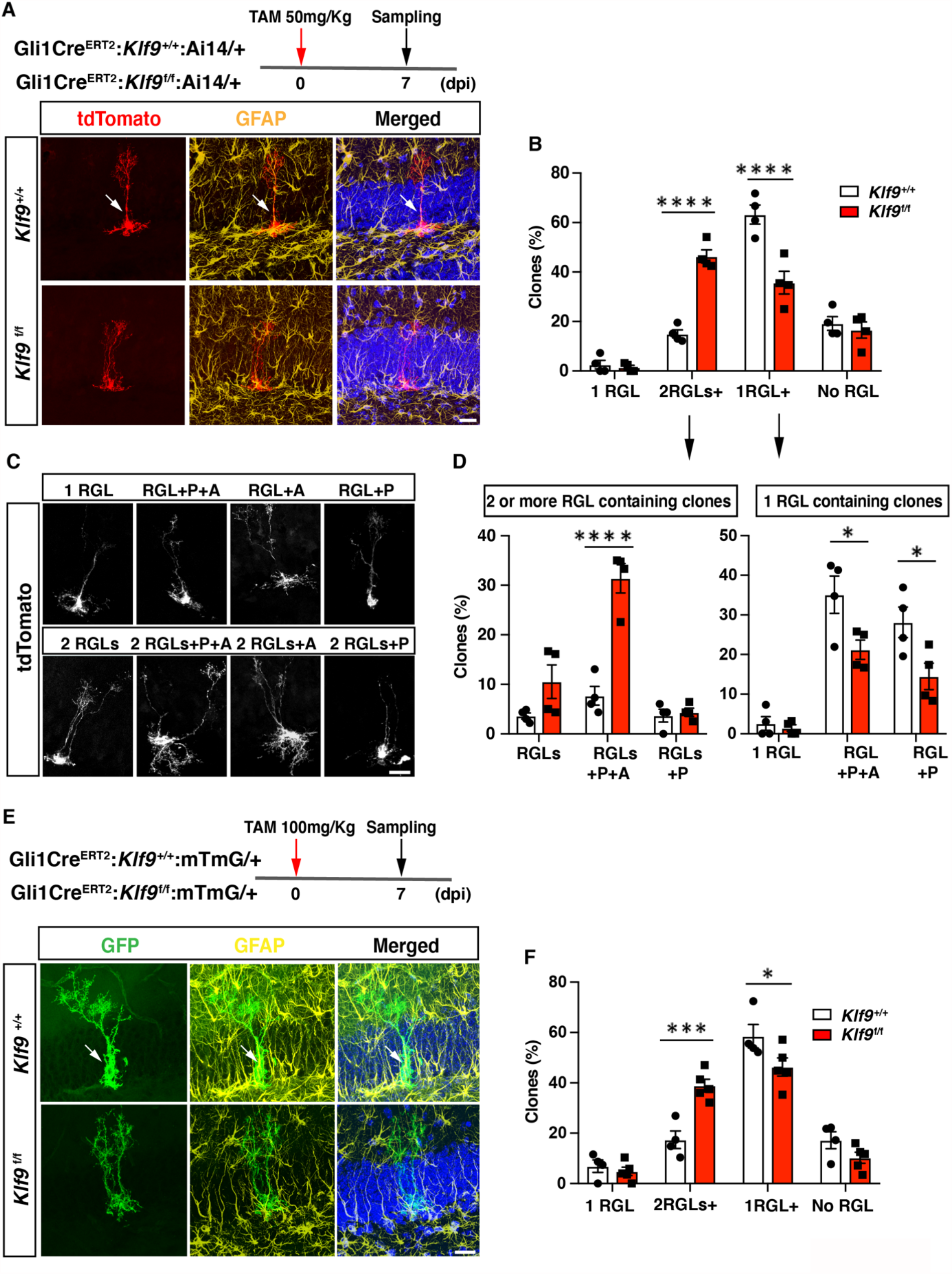
*Klf9* deletion in RGLs produces supernumerary RGL clones. (A-D) Clonal analysis of sparsely labeled Gli1+RGLs in adult Gli1Cre^ERT2^*:Klf9*^+/+or f/f^:Ai14 mice at 7dpi. (A, C) Representative images of labeled RGL clones and descendants. For example, A top: single RGL (white arrow), A bottom: 2 RGLs. Identification was based on tdTomato+ morphology and GFAP immunohistochemistry. (B) Statistical representation of clones for specified compositions for both genotypes expressed as fraction of total clones quantified. (D) Breakdown of clones into 2RGL+ (2 or more RGL containing clones and progeny) and single RGL+ clones (clones containing only 1 RGL and progeny). Loss of Klf9 in Gli1+ RGLs results in statistically significant overrepresentation of 2 or more RGL containing clones and significant reduction in “1 RGL containing clones” suggestive of *Klf9* repressing RGL expansion. n=4 mice/group. (E-F) Clonal analysis of sparsely labeled Gli1+RGLs (white arrow) in adult Gli1Cre^ERT2^*:Klf9*^+/+or f/f^:mTmG mice at 7dpi. Inducible deletion of *Klf9* in Gli1+ RGLs results in statistically significant overrepresentation of multiRGL containing clones (2 or more RGLs, 2RGL+) and a significant reduction in single RGL containing clones (1RGL+). Identification was based on GFP+morphology and GFAP immunohistochemistry. Representative images (E) and corresponding quantification in (F). n=4 and 5 mice/group. P: Progenitor, A: Astrocyte. Data are represented as mean ± SEM. * p<0.05, *** p<0.001, ****p<0.0001. Scale Bar Figures 2A, 2C and 2E 20 μm.

### *Klf9* functions as a brake on symmetric self-renewal of RGLs

To unequivocally establish clonal origin of labeled progeny and directly test the hypothesis that *Klf9* inhibits symmetric self-renewal of RGLs *in vivo*, we performed longitudinal two-photon imaging (*46*) of RGLs for up to two months and tracked symmetric and asymmetric division patterns (**Figures 3A-B; Figures S3A-B, Supplementary Movies S1-S4**). We implanted Gli1Cre^ERT2^:*Klf9*^f/f or +/+^;Ai14 mice with a hippocampal window over CA1 for long term imaging. After allowing 2 weeks for recovery from surgery, we injected mice with a single dose of tamoxifen (150mg/kg) to induce Cre recombination and tdTomato expression in Gli1+ RGLs. This resulted in sparse labeling that allowed us to image and track individual cells and their processes. Imaging sessions started 2 days post tamoxifen injection (dpi) and were repeated daily up to 6 dpi in order to locate isolated labeled RGLs which were clearly identifiable by their tufted radial process. Astrocytes were occasionally labelled but were readily distinguishable from RGLs due to their lack of polar morphology and were disregarded. Post-hoc histology analysis of morphological features and immunoreactivity for GFAP in brain sections corroborated our initial *in vivo* identification of RGLs (**Figure S3**). After 6 dpi we track individual RGLs to quantify their first division event: we revisited each previously identified, RGL-containing field-of-view every three days and compared it with previous timepoints in order to quantify the first cell division and classify it as symmetric or asymmetric. As previously described (*9*), asymmetric divisions resulted in motile daughter cells that migrated away from their progenitors within 1-2 imaging sessions (3-6 days) and exhibited shorter and less stable processes, often undergoing further divisions and differentiation (**Figure 3B**)**(Supplementary Movie S1)**. Conversely, and as shown previously (*9*), symmetric divisions resulted in the appearance of a faint radial process of a single static daughter cell that remained adjacent to its mother cell (**Figure 3B)(Supplementary Movies S2-S4)**. Over time the cell body of the daughter RGL emerged. For our analysis of cell division, we only considered the first division event from an identified RGL, disregarding subsequent divisions of the daughter cells and analysis of RGL derived lineage trees. Deletion of *Klf9* in RGLs resulted in a significantly greater number of symmetric cell divisions (39 symmetric, 38 asymmetric, 10 mice) compared to *Klf9*^+/+^ RGLs (18 symmetric, 47 asymmetric, 8 mice) (**Figure 3C**). We made sure to have a similar number of division events across both genotypes so that we were confident that the differences in the mode of division are not due to under/over sampling each experimental group (N= 8 control mice, 65 divisions, mean 8.125 divisions per mouse; 10 experimental mice, 77 divisions, mean 7.7 divisions per mouse) (**Figure 3D**). These data provide definitive evidence for *Klf9* functioning as a brake on symmetric self-renewal of RGLs in the adult hippocampus.

**Figure 3.**
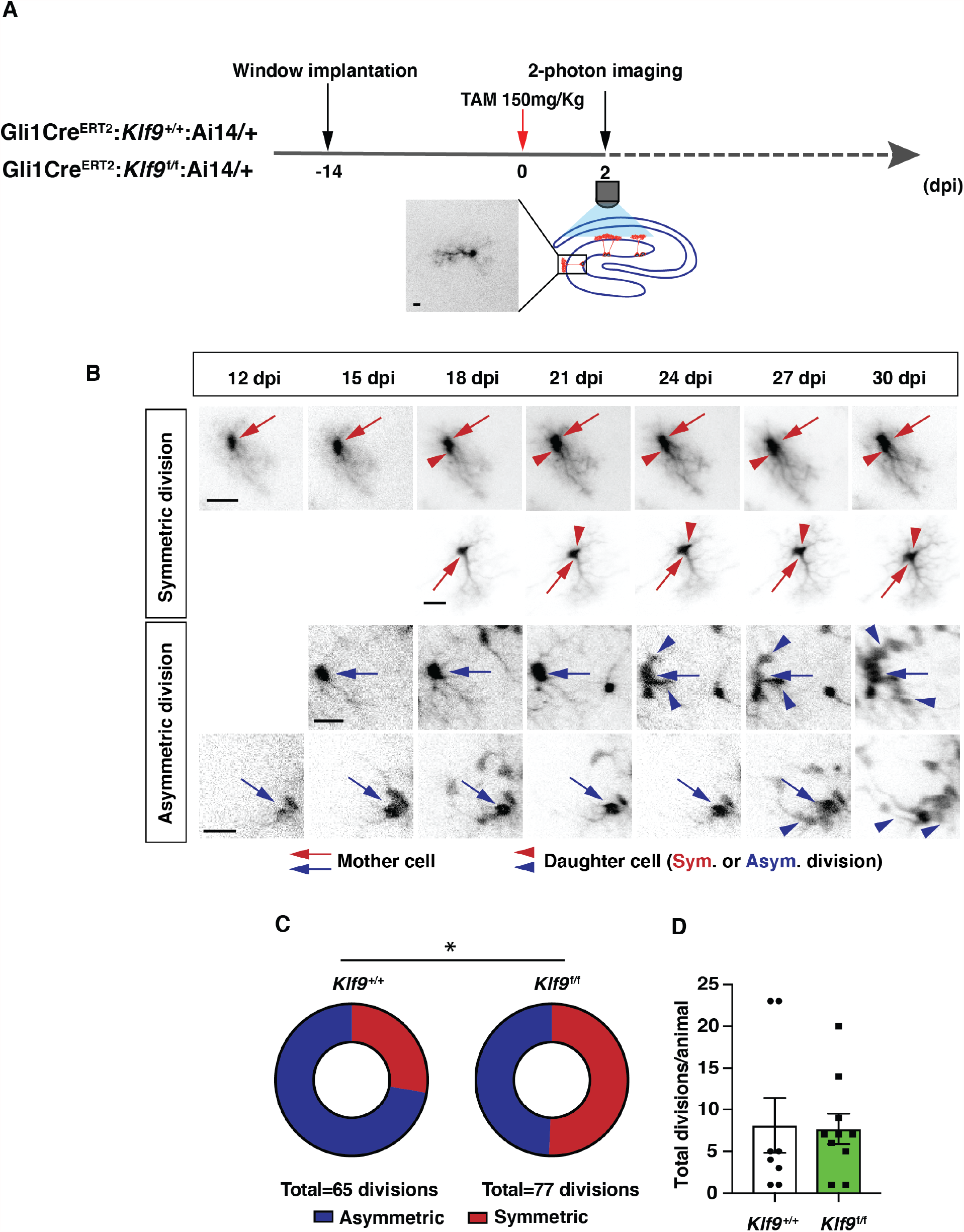
Klf9 functions as a brake on symmetric self-renewal of RGLs. (A) Diagram of experimental design for *in vivo* 2-photon imaging experiments. Inset is a high magnification image of a sparsely labeled single RGL in an adult Gli1Cre^ERT2^*:Klf9*^+/+^:Ai14 mouse. (B) Representative series of longitudinal imaging from four fields of view showing RGL symmetric and asymmetric divisions. Arrows point to mother cell and arrowheads point to daughter cells. Scale bars: 20 µm. (C) Quantification of RGL symmetric and asymmetric divisions showing an increase in symmetric divisions in Gli1Cre^ERT2^*:Klf9*^f/f^:Ai14 mice. n=8 Gli1Cre^ERT2^*:Klf9*^+/+^:Ai14 mice, 65 divisions; n=10 Gli1 Cre^ERT2^*:Klf9*^f/f^:Ai14 mice, 77 divisions. Odds of symmetric division are 2.7x higher in *Klf9*f/f, p=0.015 Likelihood-Ratio Test. (D) Similar number of divisions were recorded for each group to avoid biased assessment of division mode (n=8 and 10 mice/group). See also corresponding Supplementary Figure 3 and Supplementary Movies S1, S2, S3 and S4.

### *Klf9* regulates a genetic program of RGL activation and expansion

To understand how Klf9 regulates RGL division mode, we performed *in vivo* molecular profiling of RGLs lacking *Klf9*. We generated Gli1Cre^ERT2^:Rpl22HA^f/+^:*Klf9*^f/f or +/+^ (B6N.129-*Rpl22tm1*.*1Psam*/J mice (Ribotag) mice (*47*) to genetically restrict expression of a hemagglutinin (HA) epitope-tagged ribosomal subunit exclusively in Gli1+ RGLs (**Figure 4A**). Four days following TAM injections to induce HA expression and Klf9 recombination in a sufficient number of Gli1+RGLs and progeny arising from first division, we dissected the dentate gyrus subregion, biochemically isolated actively translated transcripts, generated cDNA libraries and performed Illumina sequencing **(Figures 4A-C)**. Analysis of the resulting data and gene ontology annotation (gGOSt, https://biit.cs.ut.ee/gprofiler/gost) of differentially expressed genes (DEGs)**(Supplementary Table 1)** broadly categorized signaling pathways and molecular programs associated with neural stem cell activation and quiescence (*32, 34, 48-52*) (**Figure 4C; Figure S4, Supplementary Tables 2 and 3**). Functional categories enriched among upregulated DEGs (276) included phospholipase activity (Pla2g7,Pla2g4e,Gpld1), mitogen growth factor signaling (Egfr, Fgfr3, Ntrk2, Lfng) and ligand-gated ion channels (Gabra1, Chrna7, Grin2C, P2rx7). Additionally, our analysis revealed elevation of metabolic programs sustaining energy production and lipogenesis through generation of Acetyl-CoA: CoA- and fatty acid-ligase activity (Acsl3, Ascl6, Acss1, Acsbg1) and oxidoreductase and aldehyde dehydrogenase activity (Acad12, Acox1, Ak1b10, Aldh3b1, Aldh4a1)(*52-55*)**(Supplementary Table 2)**. A complementary set of modules over-represented in the downregulated gene set (462 DEGs) were quiescence growth factor signaling (Bmp2, Bmp4), extra-cellular matrix binding (Itga3, Itga10, Igfbp), cell adhesion (eg: Emb, Itga3, Itga5), actin binding (Iqgap1), transcription factors (NeuroD4, Zic3) and voltage-gate potassium channel activity (Kcnj8, Kcnq1)**(Supplementary Table 3)**.

**Figure 4.**
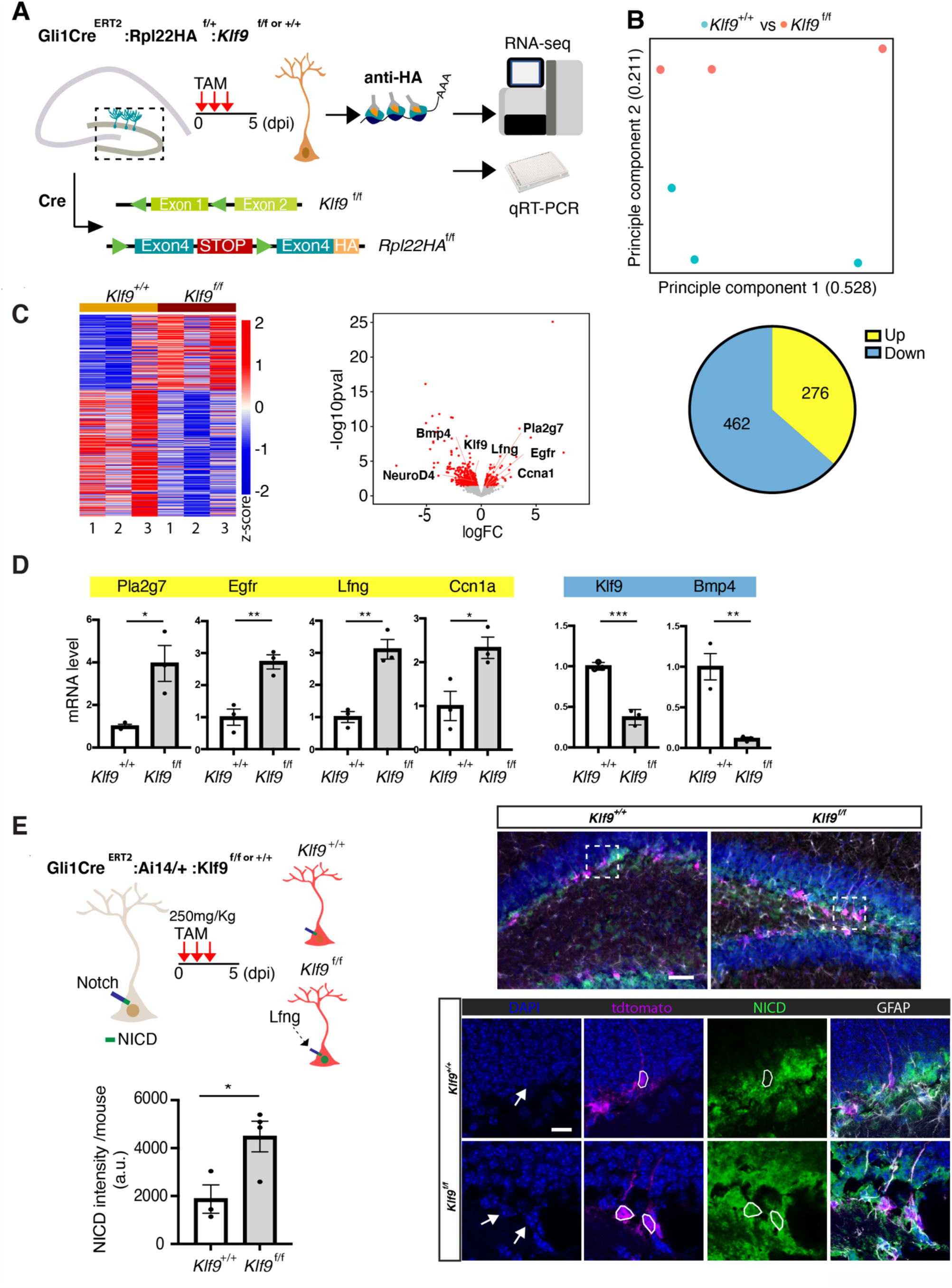
*Klf9* regulates genetic programs underlying RGL expansion. (A) Schematic of experimental workflow to biochemically isolate and sequence translated mRNAs from Gli1+RGLs (Gli1Cre^ERT2^:Rpl22HA^f/+^:*Klf9*^f/f or +/+^ mice). n=3 mice, 6 dentate gyri/sample, 3 samples per group. (B) Principal component analysis (PCA) plot of translational profiles of Gli1Cre^ERT2^ targeted *Klf9*^+/+ or f/f^ RGLs. First two principal components are shown with the corresponding fractions of variance. (C) Left: Heatmap of expression values for differentially expressed genes. Middle: Volcano plot of statistical significance (-log10 P-value) vs. magnitude of change (log2 of fold change) of gene expression. Differentially expressed genes are marked in red. Upregulated genes in *Klf9*^f/f^ RGLs are on the right and downregulated genes in *Klf9*^f/f^ RGLs are on the left. Right: Piechart of numbers of upregulated and downregulated genes in Gli1Cre^ERT2^ targeted *Klf9*^f/f^ RGLs. (D) qRT-PCR on biochemically isolated mRNAs from Gli1Cre^ERT2^:Rpl22HA^f/+^:*Klf9*^f/f or +/+^ mice validating candidate differentially expressed genes. n=3 mice, 6 dentate gyri/sample, 3 samples per group. (E) Immunostaining and quantification of NICD in RGLs of Gli1Cre^ERT2^: *Klf9*^f/f or +/+^ mice. Deletion of *Klf9* results in increased NICD levels in RGLs consistent with Lnfg dependent potentiation of Notch1 signaling (Cartoon, top left). n=3 mice/group. Data are represented as mean ± SEM. * p<0.05, ** p<0.01, *** p<0.001. Scale bar: top, 20 um; bottom, 10 um. See also corresponding Supplementary Figure 4 and Supplementary Tables 1-3.

For validation of DEGs previously linked with neural stem cell quiescence and activation (*32, 34, 49, 50, 52, 56*), we performed qRT-PCR on an independent replicate of biochemically isolated mRNAs from this population of Gli1+RGLs *in vivo*. We first confirmed downregulation of Klf9 in RGLs. Next, we validated downregulation of canonical quiescence signaling factors (Bmp4) and upregulation of genes involved in lipid metabolism (Pla2g7), cell cycle (Ccn1a), mitogen signaling (epidermal growth factor receptor, Egfr), and Notch signaling (Lunatic fringe, Lfng) **(Figure 4D)**. Consistent with Lfng-mediated potentiation of Notch1 signaling through cleavage of the Notch1 intra-cellular domain (NICD), we observed significantly elevated levels of NICD in Gli1+ RGLs lacking Klf9 (**Figure 4E**)(*51, 57*). We infer from our loss-of function data that high levels of Klf9 in RGLs induce BMP4 expression and repress gene modules specifying mitogen signaling, fatty acid oxidation, RGL differentiation and cell-cycle exit to inhibit RGL expansion.

## Discussion

Central to experience-dependent regulation of neurogenesis is the ability of RGLs to constantly balance demands for neurogenesis and astrogenesis or RGL expansion with self-preservation through regulation of quiescence. Since interpretation of the external world is dependent on integration and convergence of physiological extracellular signals upon TFs in RGLs, enriching and adverse experiences are likely to modulate the balance between transcriptional programs that regulate RGL division modes supporting amplification or asymmetric self-renewal (*22*). However, in contrast to our knowledge of TFs that regulate asymmetrical self-renewal of RGLs in the adult hippocampus (*32-36*), the identities of transcriptional regulators of symmetric self-renewal of RGLs have remained elusive. By combining conditional mouse genetics with *in vivo* clonal analysis and longitudinal two-photon imaging of RGLs, we demonstrated that Klf9 acts as a transcriptional brake on RGL activation state and expansion through inhibition of symmetric self-renewal (**Figure 5**).

**Figure 5.**
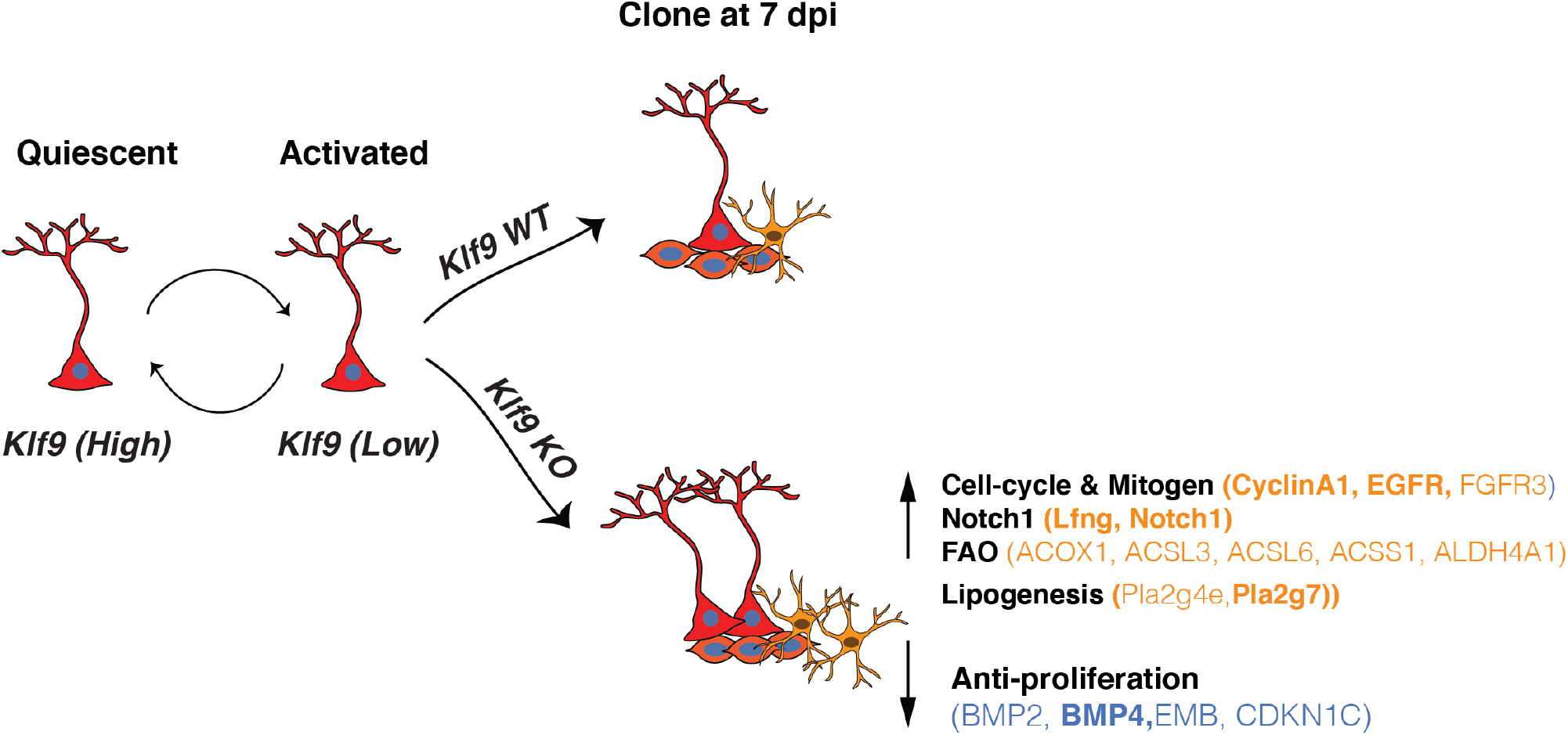
Summary schematic conveying *Klf9* functions in RGL activation and self-renewal. RGLs integrate extracellular, experiential signals to exit quiescence, the dominant state, and become activated. *Klf9* expression is elevated in quiescent RGLs. Low levels of *Klf9* in RGLs is associated with increased activation. Once activated, RGLs lacking *Klf9* are biased towards symmetric self-renewal and RGL expansion. Translational profiling of RGLs reveals how loss of *Klf9* results in downregulation of a program of quiescence and activation of genetic (mitogen, notch) and metabolic (fatty acid oxidation and lipid signaling) programs underlying RGL symmetric self-renewal. Candidate differentially expressed upregulated (orange) and downregulated genes (blue) in RGLs following *Klf9* deletion are shown here. Genes in bold indicates validation by qRT-PCR.

That Klf9 expression is higher in non-dividing RGLs than in activated RGLs is consistent with gene expression profiling of quiescent adult hippocampal RGLs (*42, 52*)(Jaeger and Jessberger, personal communication) and other quiescent somatic stem cells such as satellite cells (*58*) and neural stem cells in the subventricular zone (*49, 59, 60*). Loss of Klf9 in Gli1+ RGLs resulted in increased RGL activation. Based on our clonal analysis of RGL output and in vivo translational profiling, we think that this increased RGL activation reflects maintenance of an activated or cycling state (also discussed later) to support increased symmetric self-renewal (*6*).

Our current knowledge of TFs that regulate symmetric self-renewal in the adult hippocampus can only be extrapolated from studies on hippocampal development (*61*). Clonal analysis of Gli1-targeted RGLs revealed multi RGL containing clones with progeny. This potentially reflects competition between TFs that dictate balance between symmetric and asymmetric divisions, compensation by downstream effectors of Klf9 or constraints on RGL expansion imposed by availability of niche factors. Such compensatory mechanisms may also explain why constitutive deletion of *Klf9* does not overtly affect size of the dentate gyrus (*39*).

Studies on adult hippocampal neural stem and progenitor cells have relied on assays that induce quiescence and activation *in vitro* (*52*),unbiased single cell profiling of neurogenesis (*50, 51*) or FACS sorting of neural stem and progenitor cells *in vivo* (*34*). Because asymmetric self-renewal is the predominant mode of division, it is most certainly the case that the RGL activation profile inferred from these studies is biased towards asymmetric, rather than symmetric, self-renewal. In contrast, our *in vivo* translational profiling of long-term self-renewing Gli1+RGL population following cell-autonomous deletion of Klf9 allowed us to infer how changes in gene expression relate to RGL symmetric division mode and create an exploratory resource for the neural stem cell community. While ribosomal profiling does not allow us to isolate transcripts from single RGLs, it offers other advantages such as minimizing stress response associated with cell dissociation (*62*). Since Gli1CreERT2 specifically targets RGLs and astrocytes (but not progenitors) and we performed biochemical profiling at 4 days post recombination when we first observe RGL derived progeny, our analysis largely reflects changes in the RGL population, progeny arising from first division and astrocytes. That we observe an enrichment of genes expressed exclusively in RGLs permits us to link gene expression with changes in RGL numbers driven by division mode. Analysis of Klf9 levels by qRT-PCR suggests greater than 50% recombination efficiency of Klf9 in targeted cell populations. Our genome-wide expression analysis suggests that Klf9 functions as an activator or repressor depending on cellular context, although repression appears to be the dominant mode of gene regulation (*63, 64*). Validation of specific DEGs in biochemically isolated transcripts from RGLs suggests that Klf9 may activate BMP4 expression in RGLs to suppress activation *in vivo* (*48*). Additionally, Klf9 suppresses RGL proliferation through repression of mitogen signaling receptor tyrosine kinases (EGFR), lipidogenesis (Pla2g7) and cell-cycle (CyclinA1). Pla2g7, interestingly, is expressed only in RGLs and astrocytes in the DG (*50, 51*) and as such may represent a novel marker of activated RGLs. Given the dual roles of Notch signaling in regulation of active and quiescent RGLs (*65*), we validated that Lfng is significantly upregulated in RGLs lacking Klf9. Lfng is exclusively expressed in RGLs, promotes Notch1 signaling through glycosylation of Notch1 and generation of NICD following ligand binding, and dictates RGL activation in a ligand dependent manner (*66*). Genetic overexpression of Lfng in T cell progenitors sustained Notch1 mediated self-renewal and clonal expansion at expense of differentiation (*67*). Consistent with Lfng upregulation in RGLs, we observed elevated levels of NICD in Gli1+ RGLs lacking Klf9 indicative of enhanced Notch1 signaling.

Bioinformatics analysis of our data identified enhanced fatty acid β-oxidation (FAO), a substrate for energy production and lipogenesis as a metabolic program recruited to sustain RGL expansion (**Figure 5**). In fact, lineage tracing studies on embryonic neocortical neural stem cells has demonstrated a role for FAO in maintenance of neural stem cell identity and proliferation(*53*). Specifically, inhibition of Tmlhe (a carnitine biosynthesis enzyme) and carnitine-dependent long-chain fatty acid β-oxidation (carnitine palmitoyltransferase I, CPT1, which catalyzes the rate-limiting reaction in this process) resulted in a marked increase in symmetric differentiating divisions at expense of both symmetric and asymmetric self-renewal of neural stem cells (*55*). Inhibition of FAO prevented hematopoietic stem cell maintenance and promoted symmetric differentiating divisions of hematopoietic stem cells (*68*). Deletion of Cpt1a (and inhibition of FAO) in adult hippocampal neural stem cell and progenitors impaired expansion and reduced numbers of RGLs, although it could not be determined if this was due to death and/or inhibition of symmetric self-renewal of RGLs (*54*).

How does Klf9 function as a brake on RGL symmetric self-renewal? We propose that Klf9 co-represses a suite of genes associated with maintenance of RGLs in symmetric division mode. Pioneering studies have implicated Notch signaling in sustaining symmetric divisions of neuroepithelial cells (*69*), expansion of putative neural stem cells and progenitors (*70*) and maintenance of radial glial cell like identity through inhibition of differentiation and cell-cycle exit (*71, 72*). Importantly, genetic gain-of-function of Notch1 signaling in RGLs in the adult DG maintains RGLs at the expense of hippocampal neurogenesis (*73*). Klf9 may also directly suppress a pro-neurogenic program in RGLs (for eg: NeuroD4, downregulated DEG, **Supplementary Table 3**)(*74*) or indirectly via competitive interactions with TFs that regulate RGL asymmetric self-renewal. Taken together, loss of Klf9 in RGLs drives expansion through enhanced mitogen and cell-cycle signaling (*75*), prevention of RGL differentiation, and elevation of lipogenic and FAO metabolic programs (**Figure 5**).

Our findings stimulate discussion on how experiential signals regulate RGL activation and expansion. To date, GABA(A) R signaling and PTEN signaling (by inhibiting PI3K-Akt pathway) have been shown to promote quiescence and suppress RGL amplifying divisions (*5, 25*). It is plausible that Klf9 participates in these signaling pathways as a downstream actuator. *Klf9* expression is reduced in neural stem cells lacking FoxO3 (*60*). Thus, Akt dependent regulation of neural stem cell activation through inactivation of FoxO3 (*31*) may require *Klf9* downregulation. Since some of the identified Klf9 target genes are also regulated by other TFs [(eg: inhibition of EGFR and cyclinA1 by Notch2 (*34*), activation of Pla2g7 by FoxO3 (*60*)], we infer that these factors do not compensate each other, but instead, confer flexible integration of diverse physiological signals in RGLs to regulate activation. Inhibition of pulsatile glucocorticoid receptor signaling has also been shown to promote RGL quiescence (*24*). Because Klf9 expression is regulated by steroid hormone signaling and neural activity (*39, 76, 77*) and Klf9 represses gene expression through recruitment of a mSin3A co-repressor complex (*78*), Klf9 may support an epigenetic mechanism for reversible, experiential regulation of NSC decision making.

Our genome-wide dataset serves as a general exploratory community resource in several ways. First, it catalyzes further enquiry into mechanisms underlying neural stem cell quiescence and expansion. By way of example, candidate genes such as the cell adhesion molecule Embigin (downregulated DEG) regulates quiescence of hematopoietic stem/progenitor cells (*79*) whereas the alpha7 nicotinic receptor (upregulated DEG), ChrnA7, has been shown to be required for maintaining RGL numbers (*80*). Second, numerous genes identified in our blueprint are implicated in driving tumorigenesis and as such may guide differentiation-based strategies to block tumor proliferation (*81*). Third, our work motivates assessment of how Klf9 may link extracellular, physiological signals with genetic and metabolic programs in RGLs. Fourth, our findings may guide investigation of functional significance of Klf9 enrichment in other quiescent neural (SVZ)(*49, 59, 60*) and somatic stem cell populations (*58*).

Our study enables a more holistic assessment of how competing transcriptional programs in RGLs mediate decision making by including regulators of symmetric and asymmetric self-renewal. A deeper understanding of Klf9-dependent regulation of RGL homeostasis may guide genetic and metabolic strategies to replenish the RGL reservoir and restore neurogenesis following injury or expand the NSC pool in anticipation of future neurogenic demands to support hippocampal dependent memory processing and emotional regulation (*19, 20, 38*).

## Materials and Methods

Animals were handled and experiments were conducted in accordance with procedures approved by the Institutional Animal Care and Use Committee at the Massachusetts General Hospital and Albert Einstein College of Medicine in accordance with NIH guidelines. Mice were housed three to four per cage in a 12 hr (7:00 a.m. to 7:00 p.m.) light/dark colony room at 22°C–24°C with ad libitum access to food and water.

### Mouse lines

The following mouse lines were obtained from Jackson Labs: *Klf9*-*lacZ* knock-in (Stock No: 012909), Gli1Cre^ERT2^ (Stock No: 007913), Ai14 (Stock No: 007908), mT/mG (Stock No: 007676), B6N.129- *Rpl22*^*tm1*.*1Psam*^/J (RiboTag)(Stock No: 011029), POMC-Cre (Stock No. 010714). Sox1tTA transgenic mice(*44*) were obtained from Dr. Robert Blelloch (University of California, San Fransisco). *Klf9*^*LacZ/LacZ*^ mice were obtained from Dr. Yoshiaki Fujii-Kuriyama (University of Tsukuba and is also available from Jackson Labs, Stock No: 012909). tetO Kf9/Klf9 knock-in mice were generated by us previously (*38*). Nestin GFP mice (*40*) were obtained from Dr. David Scadden at MGH.

*Klf9* conditional knockout mice were generated through homologous gene targeting using C57BL/6 ES cells by Cyagen. F0s were bred with C57BL/6J mice to generate F1s with germline transmission and mice were backcrossed with C57BL/6J mice for 5+ generations. A set of primers (Forward: GGTAGTCAAATGGCGCAGCTTTT; Reverse: CCATCCATTCCTTCATCAGTCTCC) was used to genotype *Klf9*^*+/+ or f/f*^ mice to amplify 363 bp mutant band and 240 bp wildtype band. Gli1Cre^ERT2^: *Klf9*^*+/+ or f/f*^ Ai14, Gli1Cre^ERT2^: *Klf9*^*+/+ or f/f*^:mT/mG+/-, were generated by crossing Gli1Cre^ERT2^ mice with mT/mG or Ai14 and *Klf9*^*+/+ or f/f*^ mice.

### BrdU administration

For analysis of cell proliferation in dentate gyrus, mice were injected with BrdU (200 mg/kg body weight, i.p.) and sampled 2 hours later. For analysis of long-term retaining cells in dentate gyrus, mice were given daily injection of BrdU (25 mg/Kg body weight, i.p.) for 14 days and sampled 24 hours after the last injection.

### Tamoxifen administration

Tamoxifen (20 mg/ml, Sigma, T5648) was freshly prepared in a 10% ethanol of corn oil (Sigma C8267). For population analysis, a dose of 150 mg/kg or 250 mg/Kg was intraperitoneally injected into 8 weeks old male and female mice (**Figure 1F**). For clonal analysis, a dose of 50mg/Kg and 100mg/Kg were used in reporter lines of Ai14 and mT/mG respectively (**Figure 2A and Figure 2E**). Mice were sampled 7 or 28 days post-tamoxifen injection. For two-photon imaging (**Figure 3A**), one dose of 150 mg/kg Tamoxifen was given 2 days prior to *in vivo* imaging. For ribosomal profiling, a dose of 375 mg/kg body weight was intraperitoneally injected into 2-3 months mice every 12 hours for 3 times. Mice were sampled 4 days after the last injection (**Figure 4A**).

### Tissue processing and immunostaining

35 μm cryosections obtained from perfused tissue were stored in PBS with 0.01% sodium azide at 4°C. For immunostaining, floating sections were washed in PBS, blocked in PBS containing 0.3% Triton X-100 and 10% normal donkey serum and incubated with primary antibody overnight at 4 °C overnight (Rockland, rabbit anti RFP, 1:500; Millipore, chicken anti-GFAP, 1:2000; goat anti-GFP, Novus, 1:500; Santa Cruz, sc-8066, Goat anti-DCX, 1:500). The Mcm2 (BD Biosciences, mouse anti-Mcm2; 1:500), GFP (Abcam, Chicken anti-GFP, 1:2000), LacZ (Promega, Mouse anti-beta Galactosidase, 1:2000) and Nestin (Aves lab, chicken anti-Nestin, 1:400) antigens were retrieved by incubating brain sections in Citric buffer in pressure cooker (Aprum, 2100 retriever) for 20 min, followed by 60 min cooling to room temperature. BrdU antigen was retrieved by incubating brain sections in 2N HCl for 30 min at 37°C following 15 mins fixation in 4% PFA on previously processed fluorescent signal. On the next day, sections were rinsed three times for 10 min in PBS and incubated for 90 min with Fluorescent-label-coupled secondary antibody (Jackson ImmunoResearch, 1:500). Sections were rinsed three times for 10 min each in PBS before mounting onto glass slides (if applicable) and coverslipped with mounting media containing DAPI (Fluoromount with DAPI, Southern Biotech). NICD (rabbit anti-cleaved Notch1, Assay Biotech Cat# L0119 RRID:AB_10687460 at 1:100) immunostaining was performed as described (*66*).

### *Klf9 in situ* hybridization

We used a transgenic mouse line that expresses GFP under the control of the Nestin promoter to label the cell bodies(*40*). Mice were sacrificed 2 hours after a single BrdU injection (200 mg/Kg). *Klf9* expression was detected by florescent in situ hybridization (FISH) using a *Klf9* antisense probe complementary to Exon 1 (530-1035bp) of *Klf9* mRNA. Briefly, in situ hybridization (ISH) was performed using dioxygenin-labeled riboprobes on 35 μm cryosections generated from perfused tissue as described (*38*). Premixed RNA labeling nucleotide mixes containing digoxigenin-labeled UTP (Roche Molecular Biochemicals) were used to generate RNA riboprobes. *Klf9* null mice were used as a negative control and to validate riboprobe specificity. Riboprobes were purified on G-50 Microspin columns (GE Healthcare). Probe concentration was confirmed by Nanodrop prior to the addition of formamide. Sections were mounted on charged glass (Superfrost Plus) slides and postfixed for in 4% paraformaldehyde (PFA). Sections were then washed in DEPC-treated PBS, treated with proteinase K (40 μg/ml final), washed again in DEPC-treated PBS, and then acetylated. Following prehybridization, sections were incubated with riboprobe overnight at 60°C, washed in decreasing concentrations of SSC buffer, and immunological detection was carried out with anti-DIG peroxidase antibody (Roche) at 4°C overnight and were visualized using Cy3-conjugated Tyramide Signal Amplification system (Perkin-Elmer) at room temperature. *In situ* hybridization was followed by immunostaining for GFP (Goat anti-GFP, Novus, 1:500), and BrdU (Rat anti-BrdU, Biorad, 1:500) incubated at 4°C overnight and followed by incubation of 488- and Cy5-conjugated secondary antibodies (Jackson ImmunoResearch, 1:500) for 2 hours at room temperature. *Klf9 in situ* hybridization was performed on POMC-Cre:*Klf9*^+/+ and f/f^ mice using *Klf9* exon1 probe to validate the Klf9 conditional knockout mice. Immunological detection was carried out with anti-DIG antibody conjugated with alkaline phosphatase (Roche). Color reaction was conducted with NBT/BCIP. *Klf9* null mice were used as a negative control.

### Images acquisition and analysis

Images were obtained from one set of brain sections (6 sets generated for each brain) for each immunostaining experiment (set of antigens). Stained sections were imaged at 20X or 40x on a Nikon A1R Si confocal laser, a TiE inverted research microscope or a Leica SP8 confocal microscope. All of analysis were performed by an experimenter blind to group identity.

*LacZ intensity quantification*. We used mice carrying a LacZ allele knocked into the endogenous *Klf9* allele (*Klf9*^*Lac/LacZ or +/LacZ*^ mice)(*39*). *Klf9*^*Lac/LacZ or +/LacZ*^ mice were crossed with Nestin GFP mice in order to map *LacZ* expression levels in quiescent RGLs (GFP+MCM2-with radial process), activated RGLs (GFP+MCM2+ with radial process), and NPCs (GFP+MCM2+ without radial processes). The distinction between RGLs and NPCs was determined through morphological analysis. Images (1024 resolution) were acquired as 7 Z-stacks with a step size of 1 μm. 2-4 stacks of images from each mouse were selected for further quantification. Since the *LacZ* gene had been knocked into the endogenous *Klf*9 locus, mean intensity of *Lac*Z expression, assessed by fluorescent signal with LacZ immunostaining using ImageJ software in each GFP+ cell body, was used as a surrogate for *Klf9* expression in *Klf9*^*+/LacZ*^ mice. Mean background intensity was obtained from *LacZ* negative regions being divided from the calculations in the same section. *Klf9* FISH signal quantification. Images (2048 resolution) were acquired by a Leica SP8 confocal microscope as 30 Z-stacks with a step size of 0.5 μm. Representative images were generated by exporting stacked confocal images at full resolution for three-dimensional visualization using Imaris. The distinction between neural progenitor cells (NPCs) and neural stem cells (NSCs) was determined through morphological analysis with GFP staining. Activated RGLs were differentiated from quiescent RGLs through BrdU antibody staining (cell proliferation markers). Analysis and quantification of *Klf9* signal intensity in each GFP+ cell body was conducted using automatic counting with ImageJ software. Images were converted into 1-bit images. Then *Klf9* puncta were counted within GFP+ cell body boundaries through particle analysis allowing for number and average size of puncta to be recorded. Klf9 intensity was calculated as puncta number multiplied by average puncta size. Background was subtracted from the calculations. Klf9 null mice crossed with Nestin GFP mice were used as a negative control.

### Clonal lineage analysis

Clonal analysis was conducted with sparse labeling after optimizing dose of Tamoxifen as previously described(*5*). Ai14 and mTmG reporter mice were used to visualize the recombined cells. Serial coronal sections were generated and immunostained for GFAP, RFP or GFP antigens. Images acquisition and analysis were restricted to entire dentate gyri ∼2000 μm along the dorsal-ventral axis. RGLs were classified as cells that were located in the subgranular zone, had radial projections that extended into the granule cell layer, and were co-labeled with GFAP and RFP or GFP. Cells with GFAP labeling without radial processes but exhibiting a bushy morphology were identified as astrocytes. Recombined GFP+ or RFP+ cells without GFAP labeling in close spatial proximity to other cells were identified as neuronal progeny cells. A single cell (astrocyte or neuron) was not counted as a clone. Images (1024 resolution) were acquired using a Leica SP8 confocal microscope as 20-25 Z-stacks with a step size of 1.5 μm. Mice with less than 2 clones per hemisection on average were determined as standard for sparse labeling and were selected for clonal analysis. Except for the single RGL clone category, all the labeled cells within one clone were in close spatial proximity to each other. Clones were categorized according to the presence or absence of an RGL and the type of progeny.

### Two-photon imaging of Gli1+ *Klf9*^*+/+ or f/f*^ RGLs division modes *in vivo*

12-16 weeks old Gli1Cre ^ERT2^:*Klf9*^*+/+ or f/f*^:Ai14 mice were used for intravital 2P imaging of RGLs.

Window Implantation: We followed an established protocol to implant a cranial window over the right hemisphere of the dorsal hippocampus (*9*). Briefly, we drilled a ∼3 mm wide craniotomy, removed the underlying dura mater and aspirated the cortex and corpus callosum. A 3-mm diameter, 1.3-mm deep titanium implant, with a glass sealed to the bottom was then placed above the hippocampus. The implant and a titanium bar (29 × 3.8 × 1.3 mm) were held in place with dental cement. A titanium bar was used in order to secure the animal to the microscope stage. Mice were given a single dose of dexamethasone (1 mg/Kg, i.p.) before surgery to reduce brain swelling, and carprofen (5mg/Kg, i.p.) for inflammation and analgesic relief after surgery completion. Implanted animals were given two weeks to recover from surgery and allow any inflammation to subside.

Two-photon imaging of aRGL divisions: *In vivo* imaging was done on a custom two-photon laser scanning microscope (based on Thorlabs Bergamo) using a femtosecond-pulsed laser (Coherent Fidelity 2, 1075 nm) and a 16x water immersion objective (0.8 NA, Nikon). We imaged mice under isoflurane anesthesia (∼1% isoflurane in O2, vol/vol) and head-fixed to the microscope stage via a titanium bar implant while resting on a 37° C electrical heating pad (RWD ThermoStar). Expression of the tdTomato fluorescent label in Gli1+ RGLs was induced with a single injection of Tamoxifen (150ul/mg) two weeks after window implantation. Imaging began two days after tamoxifen injection (2 dpi) and continued every day until 6 dpi in order to locate sparse labeled RGLs. Afterwards, mice were imaged every 3 days, whenever possible and were imaged up to 60 days. Using a coordinate system, we marked locations of RGLs for recurrent imaging of the same cell. At each time point, we acquired a three-dimensional image stack of each field-of-view containing tdTomato-expressing cells and annotated their location so that the same cell could be imaged again in the following session.

Cell division classification: Cell divisions were analyzed by two different experimenters blinded to genotype. We first compiled all z-stacks into a single sum-projected image for each time point, and then we used FIJI-ImageJ to analyze the images. Only the first recorded cell division for a given clone was included in the analysis. We defined RGL symmetric division as a new RGL generated from the mother RGL, characterized by the development of a stable radial process and static behavior of cell bodies for at least 6 days after birth. We defined asymmetric division as new neural progenitor cell(s) generated from the mother RGL that exhibited shorter and less stable processes. These NPCs often began to migrate away within 1-2 imaging sessions (3-6 days).

### Ribotag isolation of mRNAs from Gli1+RGLs

We used Gli1Cre^ERT2^:Rpl22HA^f/+^:*Klf9*^f/f or +/+^ mice which enables expression of haemagglutinin (HA)-tagged ribosomal protein RPL22 (RPL22–HA) following Cre recombination in Gli1+ *Klf9*^f/f or +/+^ RGLs. RiboTag immunoprecipitation and RNA extraction were performed 4 days after last TAM injection following the original protocol with minor modifications (*47*). 6 dentate gyri from 3 mice were pooled per sample and homogenized with a dounce homogenizer in 900 ul cycloheximide-supplemented homogenization buffer. Homogenates were centrifuged and the supernatant incubated on a rotator at 4°C for 4 hours with 9ul anti-HA antibody (CST Rb anti-HA #3724, 1:100) to bind the HA-tagged ribosomes. Magnetic IgG beads (Thermo Scientific Pierce #88847) were conjugated to the antibody-ribosome complex via overnight incubation on a rotator at 4°C. RNA was isolated by RNeasy Plus Micro kit (Qiagen 74034) following the manufacturer’s protocol. Eluted RNA was stored at -80°C. For enrichment analysis, 45ul of homogenate (pre-anti-HA immunoprecipitation) was set aside after centrifugation, kept at -20°C overnight, and purified via RNeasy Micro kit as an ‘input’ sample, and used to determine NSC enrichment. RNA quantity and quality were measured with a Tape Station (Agilent) and Qubit fluorimeter (Thermo Fisher Scientific). Sequencing libraries were prepared using Ultra Low Input RNA Kit (Clontech).

### RNA-seq analysis

NGS libraries were constructed from total RNA using Clontech SMARTer v4 kit (Takara), followed by sequencing on an Illumina HiSeq 2500 instrument, resulting in 20-30 million 50 bp reads per sample. The STAR aligner (*82*)was used to map sequencing reads to transcriptome in the mouse mm9 reference genome. Read counts for individual genes were produced using the unstranded count function in HTSeq v.0.6.0 (*83*), followed by the estimation of expression values and detection of differentially expressed transcripts using EdgeR (*84*)and including only the genes with count per million reads (CPM) > 1 for one or more samples (*85*). Differentially expressed genes were defined by at least 1.2-fold change with p< 0.05. *The data was submitted to NCBI GEO database for public access and is being processed (accession number XXX)*.

### qRT-PCR

mRNA was biochemically pooled and isolated as described above for Ribosomal profiling. The first-stranded complementary DNA was generated by reverse transcription with SuperScript IV first-strand synthesis system (Thermo Fisher Scientific). For quantification of mRNA levels, aliquoted cDNA was amplified with specific primers and PowerUp SYBR Master Mix (BioRad) by CFX384 Touch Real-Time PCR detection system (BioRad). Primers were optimized and designed to hybridize with different exons. Primers are listed here (name and sequence 5’ -> 3’ are indicated).

pla2g7 F: TCAAACTGCAGGCGCTTTTC, pla2g7 R: AGTACAAACGCACGAAGACG

Egfr F: GCCATCTGGGCCAAAGATACC, Egfr: GTCTTCGCATGAATAGGCCAAT

Lfng F: AAGATGGCTGTGGAGTATGACC, Lfng R: TCACTTTGTGCTCGCTGATC

Ccn1a F: GATACCTGCTCGGGGAAAGAG, Ccn1a R: GCATTGGGGAAACTGTGTTGA

Klf9 F: AAACACGCCTCCGAAAAGAG, Klf9 R: AACTGCTTTTCCCCAGTGTG

Bmp4 F: GACCAGGTTCATTGCAGCTTTC, Bmp4 R: AAACGACCATCAGCATTCGG

Actb F: CATTGCTGACAGGATGCAGAAGG, Actb R: TGCTGGAAGGTGGACAGTGAGG

### Statistical Analysis

Statistical analysis was carried out using GraphPad Prism software. Both data collection and quantification were performed in a blinded manner. Data in figure panels reflect several independent experiments performed on different days. An estimate of variation within each group of data is indicated using standard error of the mean (SEM). Comparison of two groups was performed using two-tailed student’s unpaired t-test unless otherwise specified. Comparison of one group across time was performed using a one-way ANOVA with repeated measure. Comparison of two groups across treatment condition or time was performed using a two-way repeated measure ANOVA and main effects or interactions were followed by Bonferroni post-hoc analysis. In the text and figure legends, “n” indicates number of mice per group. Detailed statistical analyses can be found in Supplementary Table 4. For statistical analysis of DEGs, please see RNA-seq analysis section for details.

Two-photon imaging: In order to compare differences in the modes of RGL division between the two genotypes, we used the R statistical analysis software to fit a generalized linear mixed effects (LME) model to the division numbers across different mice, using genotype as a fixed effect, and including animal identity as a random effect in order to account for differences between individual animals [DivisionType∼Genotype+(1|MouseIdentity)]. p-values were calculated with a Likelihood-Ratio Test (LRT) comparing our model to a null model with no genotype information and identical random effects [DivisionType∼1+(1|MouseIdentity)].

| **Antibodies** | **Source** | **Identifier** |
| --- | --- | --- |
| **Rat anti-BrdU** | Bio-Rad | Cat# MCA2483T,<br>RRID:AB_1055584 |
| **Rabbit anti-GFAP** | Millipore | Cat# AB5804, RRID:AB_2109645 |
| **Chicken anti-Nestin** | Aves lab | Cat# NES, RRID:AB_2314882 |
| **Rabbit anti-RFP** | Rockland | Cat# 600-401-379,<br>RRID:AB_2209751 |
| **Chicken anti-GFAP** | Millipore | Cat# AB5541, RRID:AB_177521 |
| **Goat anti-GFP** | Novus | NB100-1770, RRID:AB_10128178 |
| **Goat anti-DCX** | Santa Cruz Biotechnology | Cat# sc-8066, RRID:AB_2088494 |
| **Mouse anti- beta Galactosidase** | Promega | Cat# Z3781, RRID:AB_430877 |
| **Chicken anti-GFP** | Abcam | Cat# ab13970, RRID:AB_300798 |
| **Mouse anti-Mcm2** | BD Biosciences | Cat# 610700, RRID:AB_2141952 |
| **NICD, rabbit anti-cleaved Notch1** | Assay Biotech | Cat# L0119, RRID:AB_10687460 |
| **Rabbit anti-HA** | Cell Signaling | Cat# 3724, RRID:AB_1549585 |
| **Anti-Digoxigenin Fab fragments**<br>**Antibody, POD Conjugated** | Roche | Cat# 11207733910,<br>RRID:AB_514500 |
| **Anti-Digoxigenin Fab fragments**<br>**Antibody, AP Conjugated** | Roche | Cat# 11093274910,<br>RRID:AB_514497 |
| **Alexa Fluor® 488-, Cy3-, or Cy5-**<br>**conjugated donkey secondary** | Jackson ImmunoResearch | N/A |

## Author Contributions

Conceptualization: AS and NG

Methodology: NG, KM, YS, HZ, DG, CH, JC, AZ, WRM, LPW, RS, TG, AS

Investigation: NG, KM, YS, HZ, DG, CH, JC, AZ, WRM, LPW, RS, TG, AS

Visualization: NG, KM, YS, HZ, DG, CH, JC, AZ, WRM, LPW, RS, TG, AS

Supervision: AS, TG, RS

Writing— original draft: AS

Writing— review & editing: AS, NG, TG, KM, YS, LPW, RS

## Data and materials availability

Mouse lines generated in this study are available from the Lead Contact with a completed Materials Transfer Agreement. RNA sequencing data was submitted to NCBI GEO database for public access and is being processed (accession number XXX).

## Acknowledgements

We wish to thank members of Sahay and Goncalves labs for input on this work. N.G received support from Department of Psychiatry, MGH. KM is a trainee in the Einstein Training Program in Stem Cell Research, supported by the Empire State Stem Cell Fund through New York State Department of Health Contract C34874GG. Y.S was recipient of Taiwan Ministry of Science Postdoctoral Fellowship. D.G, C.H, J.C and A.Z are recipients of HSCI summer internship fellowships. A.S. acknowledges support from US National Institutes of Health Biobehavioral Research Awards for Innovative New Scientists (BRAINS) 1-R01MH104175, NIH-NIA 1R01AG048908-01A1, NIH 1R01MH111729-01, NINDS R56NS117529, the James and Audrey Foster MGH Research Scholar Award, Ellison Medical Foundation New Scholar in Aging, Whitehall Foundation, the Inscopix Decode Award, the NARSAD Independent Investigator Award, Ellison Family Philanthropic support, Blue Guitar Fund, Harvard Neurodiscovery Center/MADRC Center Pilot Grant Award, a Alzheimer’s Association research grant, the Harvard Stem Cell Institute (HSCI) Development grant and a HSCI seed grant. J.T.G. acknowledges support from US National Institutes of Health NINDS R56NS117529 and the Whitehall Foundation. A.S thanks L. M. S. Sahay for proof reading manuscript.

## Other Supplementary Materials for this manuscript include the following

Tables S1 to S4

**Supplementary Table 1** Complete lists of differentially expressed genes (DEGs) in Gli1+RGLs following *Klf9* deletion. DEGs were defined by at least 1.2-fold change with FDR < 0.05.

**Supplementary Table 2** Gene ontology annotation (gGOSt, https://biit.cs.ut.ee/gprofiler/gost) of differentially upregulated genes in Gli1+RGLs following *Klf9* deletion.

**Supplementary Table 3** Gene ontology annotation (gGOSt, https://biit.cs.ut.ee/gprofiler/gost) of differentially downregulated genes in Gli1+RGLs following *Klf9* deletion.

**Supplementary Table 4** Complete statistical analysis

Movies S1 to S4

**Supplementary Movie S1**: Example of longitudinal imaging of asymmetric NSC divisions. Two-photon imaging across days showing two examples of asymmetric division of NSCs (red arrows).

**Supplementary Movie S2**: Example of longitudinal imaging of symmetric cell divisions. Two-photon imaging across days showing two examples of symmetric division of NSCs (blue arrows).

**Supplementary Movie S3**: 3-dimensional reconstruction of RGL cells imaged *in vivo* before undergoing symmetric division. Field of view corresponds to second row of Figure 3B at 18 dpi.

**Supplementary Movie S4**: 3-dimensional reconstruction of RGL cells imaged *in vivo* after undergoing symmetric division. Field of view corresponds to second row of Figure 3B at 30 dpi.

**Fig. S1.**
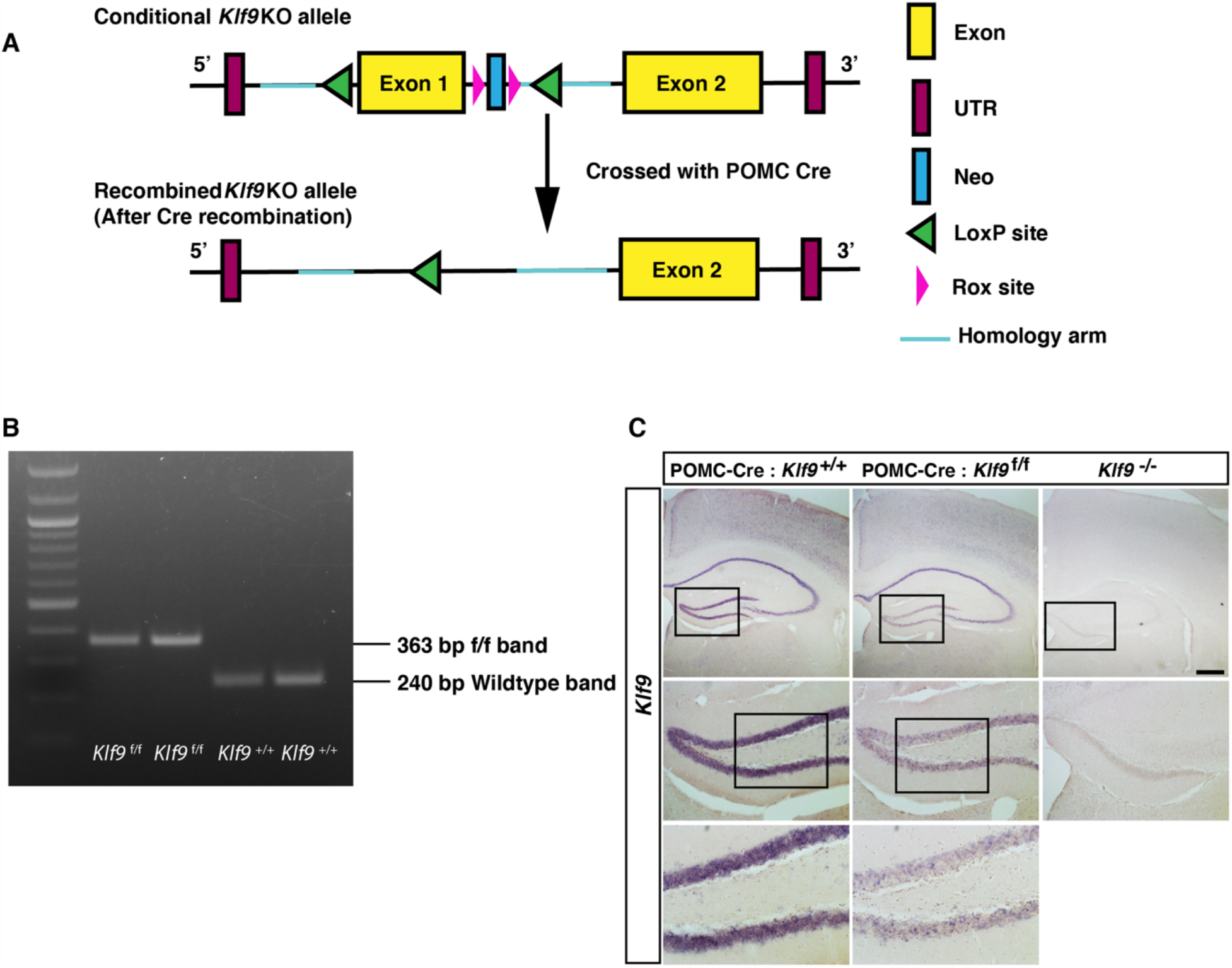
Generation and characterization of *Klf9* conditional mutant mouse line. *Related to Figure 1*. (A) Schematic of wild-type and modified *Klf9* alleles. (B) PCR on tail DNA showing expected bands conveying wild-type and conditional alleles. (C) (Left) *Klf9 in situ* hybridization on hippocampal sections from 4 months old POMC Cre: *Klf9*^+/+ or f/f^ mice showing expected salt and pepper pattern of recombination in dentate gyrus that is characteristic of POMC Cre recombination pattern in dentate gyrus. (Right) *Klf9 in situ* hybridization on hippocampal sections from adult *Klf9*^-/-^ mice conveying specificity of *Klf9* riboprobe.

**Figure S2.**
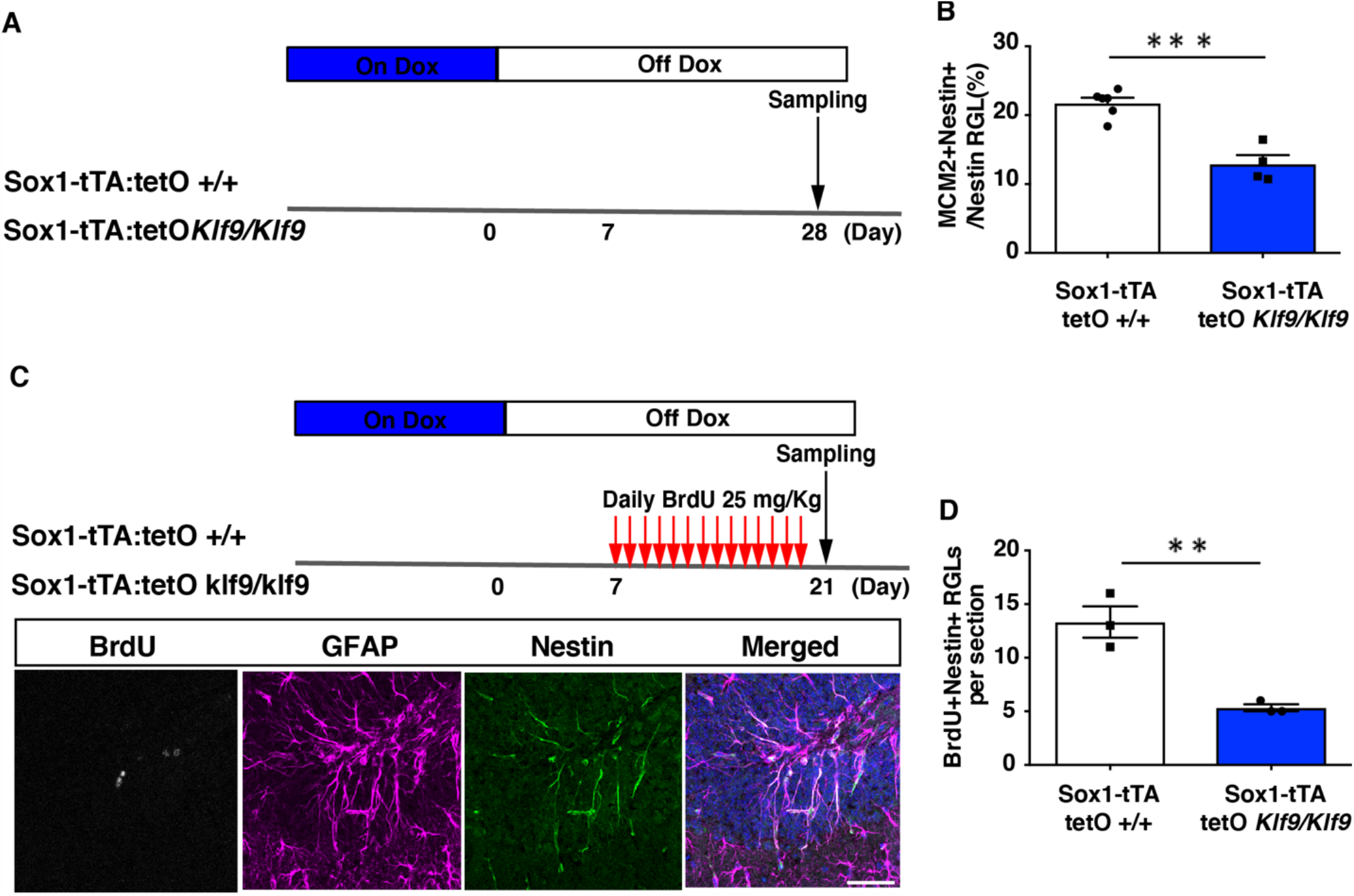
Inducible overexpression of *Klf9* in activated neural stem and progenitors promotes quiescence. *Related to Figure 1*. (A-B) Two cohorts of Adult Sox1tTA: tetO *Klf9* mice were used. *Klf9* induction in neural stem and progenitors following 3 weeks off Dox significant reduced the fraction of activated RGLs (%MCM2+Nestin+RGLS, n=6 and 4 mice/group). (C-D) A second cohort of mice was given BrdU pulses during the Off Dox window when *Klf9* is upregulated. Analysis of BrdU+Nestin+ RGLs (n=3 mice/group) revealed a significant reduction in total numbers of dividing RGLs. Representative images shown here. Unpaired t-tests, Figure S2B p=0.0003, Figure S2D p=0.005. Data are represented as mean ± SEM. ** p<0.01, *** p<0.001

**Figure S3.**
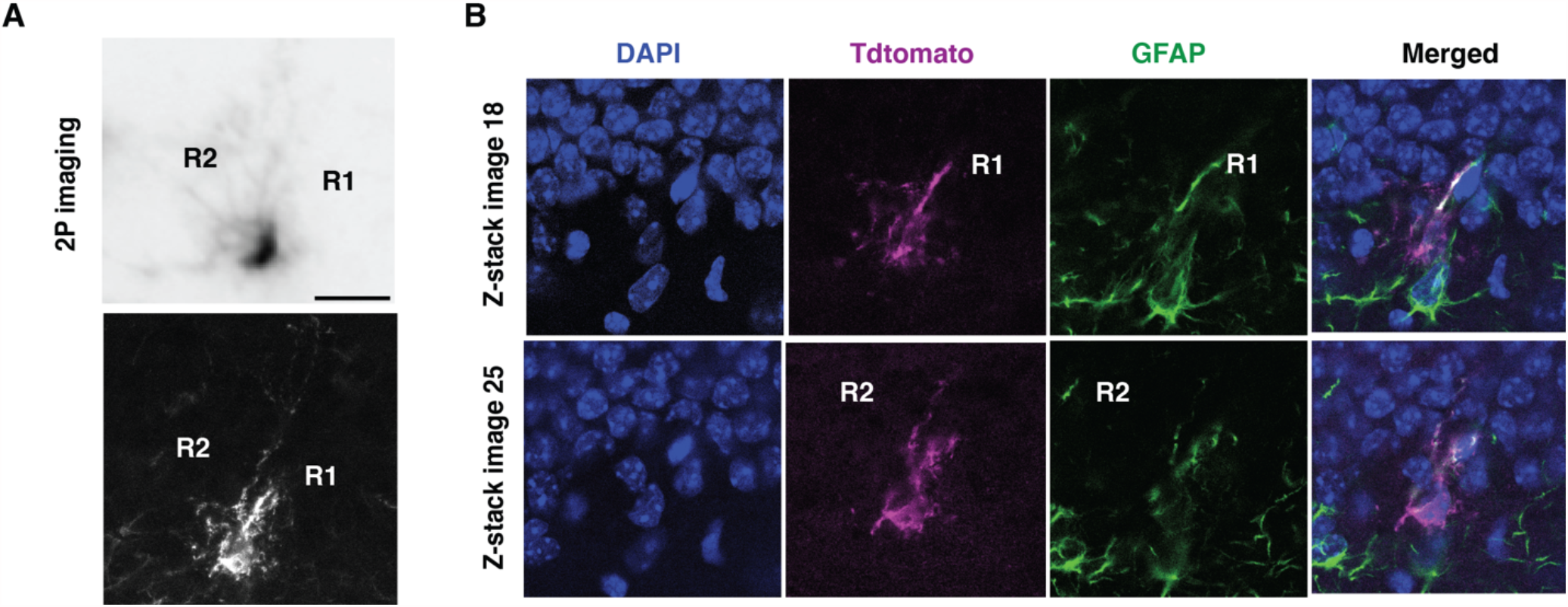
Representative images of RGL divisions captured using 2-photon imaging in vivo. *Related to Figure 3*. (A) Representative 2-photon images of RGL cells R1 and R2 *in vivo* (top) and their respective post-hoc fluorescence image (bottom). (B) Confocal immunofluorescence images of the same GFAP+/tdTomato+ cells at different depths, confirming their RGL identity. Scale bars: 20 µm.

**Figure S4.**
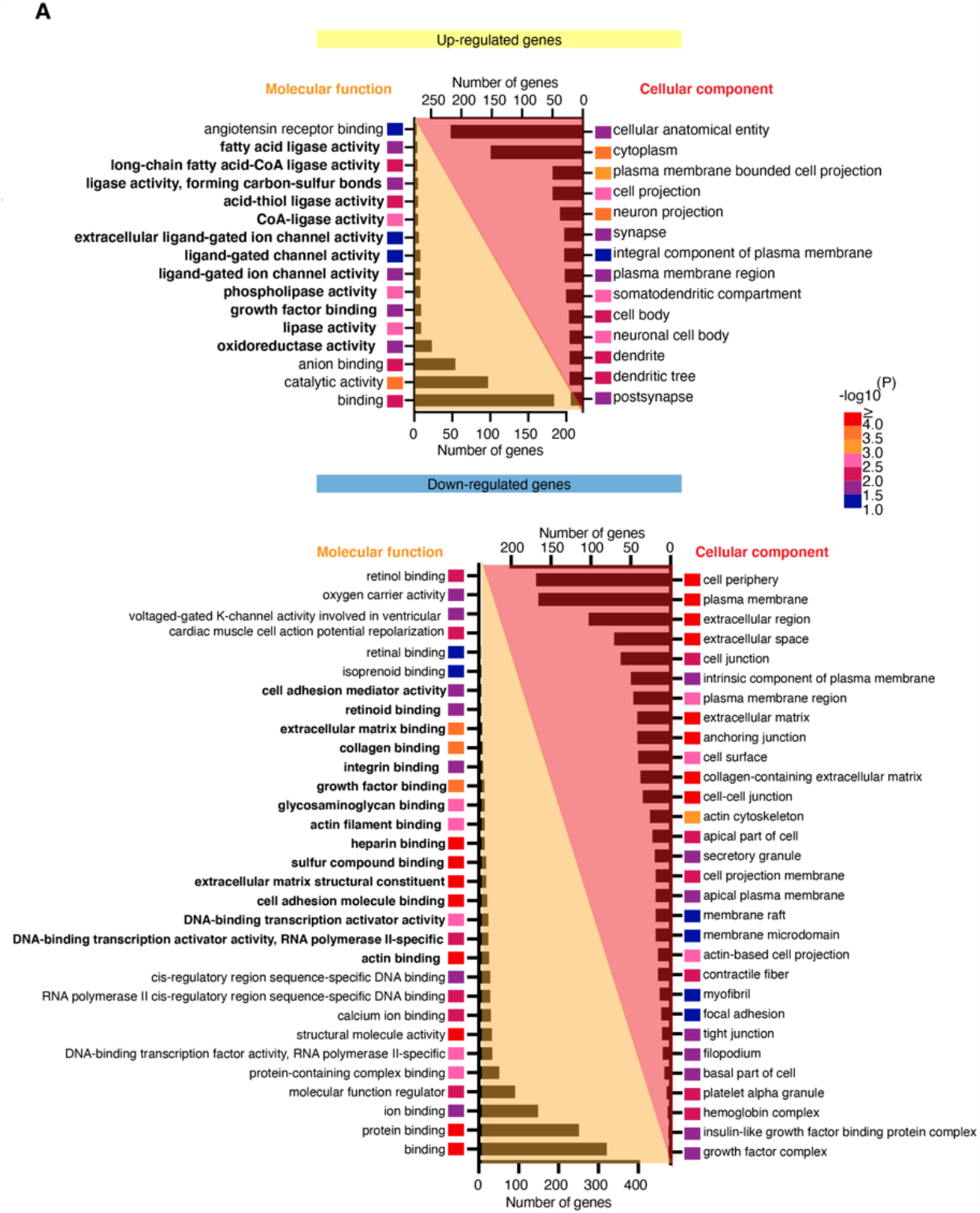
Annotation of upregulated and downregulated DEGs in Gli1+RGLs following *Klf9* deletion. *Related to Figure 4*. Gene ontology annotation (gGOSt, https://biit.cs.ut.ee/gprofiler/gost) of differentially expressed genes (DEGs) in Gli1+RGLs following *Klf9* deletion.

## References

1. B. Seri, J. M. Garcia-Verdugo, B. S. McEwen, A. Alvarez-Buylla, Astrocytes give rise to new neurons in the adult mammalian hippocampus. J Neurosci 21, 7153–7160 (2001).

2. A. D. Garcia, N. B. Doan, T. Imura, T. G. Bush, M. V. Sofroniew, GFAP-expressing progenitors are the principal source of constitutive neurogenesis in adult mouse forebrain. Nat Neurosci 7, 1233–1241 (2004).

3. S. Ahn, A. L. Joyner, In vivo analysis of quiescent adult neural stem cells responding to Sonic hedgehog. Nature 437, 894–897 (2005).

4. D. C. Lagace et al., Dynamic contribution of nestin-expressing stem cells to adult neurogenesis. J Neurosci 27, 12623–12629 (2007).

5. M. A. Bonaguidi et al., In vivo clonal analysis reveals self-renewing and multipotent adult neural stem cell characteristics. Cell 145, 1142–1155 (2011).

6. J. M. Encinas et al., Division-coupled astrocytic differentiation and age-related depletion of neural stem cells in the adult hippocampus. Cell Stem Cell 8, 566–579 (2011).

7. J. T. Goncalves, S. T. Schafer, F. H. Gage, Adult Neurogenesis in the Hippocampus: From Stem Cells to Behavior. Cell 167, 897–914 (2016).

8. J. Moss et al., Fine processes of Nestin-GFP-positive radial glia-like stem cells in the adult dentate gyrus ensheathe local synapses and vasculature. Proc Natl Acad Sci U S A 113, E2536–2545 (2016).

9. G. A. Pilz et al., Live imaging of neurogenesis in the adult mouse hippocampus. Science 359, 658–662 (2018).

10. J. Altman, G. D. Das, Autoradiographic and histological evidence of postnatal hippocampal neurogenesis in rats. J Comp Neurol 124, 319–335 (1965).

11. P. S. Eriksson et al., Neurogenesis in the adult human hippocampus. Nat Med 4, 1313–1317 (1998).

12. K. L. Spalding et al., Dynamics of hippocampal neurogenesis in adult humans. Cell 153, 1219–1227 (2013).

13. M. Boldrini et al., Human Hippocampal Neurogenesis Persists throughout Aging. Cell Stem Cell 22, 589–599 e585 (2018).

14. S. F. Sorrells et al., Human hippocampal neurogenesis drops sharply in children to undetectable levels in adults. Nature 555, 377–381 (2018).

15. E. P. Moreno-Jimenez et al., Adult hippocampal neurogenesis is abundant in neurologically healthy subjects and drops sharply in patients with Alzheimer’s disease. Nat Med 25, 554–560 (2019).

16. M. K. Tobin et al., Human Hippocampal Neurogenesis Persists in Aged Adults and Alzheimer’s Disease Patients. Cell Stem Cell 24, 974–982 e973 (2019).

17. F. H. Gage, Adult neurogenesis in mammals. Science 364, 827–828 (2019).

18. R. Knoth et al., Murine features of neurogenesis in the human hippocampus across the lifespan from 0 to 100 years. PLoS ONE 5, e8809 (2010).

19. C. Anacker, R. Hen, Adult hippocampal neurogenesis and cognitive flexibility - linking memory and mood. Nat Rev Neurosci 18, 335–346 (2017).

20. S. M. Miller, A. Sahay, Functions of adult-born neurons in hippocampal memory interference and indexing. Nat Neurosci, (2019).

21. E. C. Cope, E. Gould, Adult Neurogenesis, Glia, and the Extracellular Matrix. Cell Stem Cell 24, 690–705 (2019).

22. C. Vicidomini, N. Guo, A. Sahay, Communication, Cross Talk, and Signal Integration in the Adult Hippocampal Neurogenic Niche. Neuron 105, 220–235 (2020).

23. A. Dranovsky et al., Experience dictates stem cell fate in the adult hippocampus. Neuron 70, 908–923 (2011).

24. M. Schouten et al., Circadian glucocorticoid oscillations preserve a population of adult hippocampal neural stem cells in the aging brain. Mol Psychiatry 25, 1382–1405 (2020).

25. J. Song et al., Neuronal circuitry mechanism regulating adult quiescent neural stem-cell fate decision. Nature 489, 150–154 (2012).

26. A. Sierra et al., Neuronal hyperactivity accelerates depletion of neural stem cells and impairs hippocampal neurogenesis. Cell Stem Cell 16, 488–503 (2015).

27. A. Ibrayeva et al., Early stem cell aging in the mature brain. Cell Stem Cell, (2021).

28. K. O. Cho et al., Aberrant hippocampal neurogenesis contributes to epilepsy and associated cognitive decline. Nature communications 6, 6606 (2015).

29. L. Shahriyari, N. L. Komarova, Symmetric vs. asymmetric stem cell divisions: an adaptation against cancer? PLoS ONE 8, e76195 (2013).

30. J. Andersen et al., A transcriptional mechanism integrating inputs from extracellular signals to activate hippocampal stem cells. Neuron 83, 1085–1097 (2014).

31. N. Urban, I. M. Blomfield, F. Guillemot, Quiescence of Adult Mammalian Neural Stem Cells: A Highly Regulated Rest. Neuron 104, 834–848 (2019).

32. S. Mukherjee, R. Brulet, L. Zhang, J. Hsieh, REST regulation of gene networks in adult neural stem cells. Nature communications 7, 13360 (2016).

33. K. M. Jones et al., CHD7 maintains neural stem cell quiescence and prevents premature stem cell depletion in the adult hippocampus. Stem Cells 33, 196–210 (2015).

34. R. Zhang et al., Id4 Downstream of Notch2 Maintains Neural Stem Cell Quiescence in the Adult Hippocampus. Cell reports 28, 1485–1498 e1486 (2019).

35. O. Ehm et al., RBPJkappa-dependent signaling is essential for long-term maintenance of neural stem cells in the adult hippocampus. J Neurosci 30, 13794–13807 (2010).

36. I. Imayoshi, M. Sakamoto, M. Yamaguchi, K. Mori, R. Kageyama, Essential roles of Notch signaling in maintenance of neural stem cells in developing and adult brains. J Neurosci 30, 3489–3498 (2010).

37. D. L. Moore et al., KLF family members regulate intrinsic axon regeneration ability. Science 326, 298–301 (2009).

38. K. M. McAvoy et al., Modulating Neuronal Competition Dynamics in the Dentate Gyrus to Rejuvenate Aging Memory Circuits. Neuron 91, 1356–1373 (2016).

39. K. N. Scobie et al., Kruppel-like factor 9 is necessary for late-phase neuronal maturation in the developing dentate gyrus and during adult hippocampal neurogenesis. J Neurosci 29, 9875–9887 (2009).

40. J. L. Mignone, V. Kukekov, A. S. Chiang, D. Steindler, G. Enikolopov, Neural stem and progenitor cells in nestin-GFP transgenic mice. J Comp Neurol 469, 311–324 (2004).

41. T. J. McHugh et al., Dentate Gyrus NMDA Receptors Mediate Rapid Pattern Separation in the Hippocampal Network. Science 317, 94–99 (2007).

42. S. Bottes et al., Long-term self-renewing stem cells in the adult mouse hippocampus identified by intravital imaging. Nat Neurosci, (2020).

43. L. Madisen et al., A robust and high-throughput Cre reporting and characterization system for the whole mouse brain. Nat Neurosci 13, 133–140 (2010).

44. M. Venere et al., Sox1 marks an activated neural stem/progenitor cell in the hippocampus. Development 139, 3938–3949 (2012).

45. M. D. Muzumdar, B. Tasic, K. Miyamichi, L. Li, L. Luo, A global double-fluorescent Cre reporter mouse. Genesis 45, 593–605 (2007).

46. J. T. Goncalves et al., In vivo imaging of dendritic pruning in dentate granule cells. Nat Neurosci 19, 788–791 (2016).

47. E. Sanz et al., Cell-type-specific isolation of ribosome-associated mRNA from complex tissues. Proc Natl Acad Sci U S A 106, 13939–13944 (2009).

48. H. Mira et al., Signaling through BMPR-IA regulates quiescence and long-term activity of neural stem cells in the adult hippocampus. Cell Stem Cell 7, 78–89 (2010).

49. P. Codega et al., Prospective identification and purification of quiescent adult neural stem cells from their in vivo niche. Neuron 82, 545–559 (2014).

50. J. Shin et al., Single-Cell RNA-Seq with Waterfall Reveals Molecular Cascades underlying Adult Neurogenesis. Cell Stem Cell 17, 360–372 (2015).

51. H. Hochgerner, A. Zeisel, P. Lonnerberg, S. Linnarsson, Conserved properties of dentate gyrus neurogenesis across postnatal development revealed by single-cell RNA sequencing. Nat Neurosci 21, 290–299 (2018).

52. M. Knobloch et al., Metabolic control of adult neural stem cell activity by Fasn-dependent lipogenesis. Nature 493, 226–230 (2013).

53. T. Namba, J. Nardelli, P. Gressens, W. B. Huttner, Metabolic Regulation of Neocortical Expansion in Development and Evolution. Neuron, (2020).

54. M. Knobloch et al., A Fatty Acid Oxidation-Dependent Metabolic Shift Regulates Adult Neural Stem Cell Activity. Cell reports 20, 2144–2155 (2017).

55. Z. Xie, A. Jones, J. T. Deeney, S. K. Hur, V. A. Bankaitis, Inborn Errors of Long-Chain Fatty Acid beta-Oxidation Link Neural Stem Cell Self-Renewal to Autism. Cell reports 14, 991–999 (2016).

56. A. Baser et al., Onset of differentiation is post-transcriptionally controlled in adult neural stem cells. Nature 566, 100–104 (2019).

57. Z. Zhao, H. Wu, An Invasive Method for the Activation of the Mouse Dentate Gyrus by Highfrequency Stimulation. Journal of visualized experiments : JoVE, (2018).

58. G. Pallafacchina et al., An adult tissue-specific stem cell in its niche: a gene profiling analysis of in vivo quiescent and activated muscle satellite cells. Stem Cell Res 4, 77–91 (2010).

59. L. Morizur et al., Distinct Molecular Signatures of Quiescent and Activated Adult Neural Stem Cells Reveal Specific Interactions with Their Microenvironment. Stem cell reports 11, 565–577 (2018).

60. V. M. Renault et al., FoxO3 regulates neural stem cell homeostasis. Cell Stem Cell 5, 527–539 (2009).

61. H. Noguchi, J. G. Castillo, K. Nakashima, S. J. Pleasure, Suppressor of fused controls perinatal expansion and quiescence of future dentate adult neural stem cells. eLife 8, (2019).

62. L. Machado et al., Tissue damage induces a conserved stress response that initiates quiescent muscle stem cell activation. Cell Stem Cell, (2021).

63. J. R. Knoedler, A. Subramani, R. J. Denver, The Kruppel-like factor 9 cistrome in mouse hippocampal neurons reveals predominant transcriptional repression via proximal promoter binding. BMC Genomics 18, 299 (2017).

64. M. Ying et al., Kruppel-like factor-9 (KLF9) inhibits glioblastoma stemness through global transcription repression and integrin alpha6 inhibition. J Biol Chem 289, 32742–32756 (2014).

65. R. Sueda, R. Kageyama, Regulation of active and quiescent somatic stem cells by Notch signaling. Dev Growth Differ, (2019).

66. F. Semerci et al., Lunatic fringe-mediated Notch signaling regulates adult hippocampal neural stem cell maintenance. eLife 6, (2017).

67. J. S. Yuan et al., Lunatic Fringe prolongs Delta/Notch-induced self-renewal of committed alphabeta T-cell progenitors. Blood 117, 1184–1195 (2011).

68. K. Ito et al., A PML-PPAR-delta pathway for fatty acid oxidation regulates hematopoietic stem cell maintenance. Nat Med 18, 1350–1358 (2012).

69. B. Egger, K. S. Gold, A. H. Brand, Notch regulates the switch from symmetric to asymmetric neural stem cell division in the Drosophila optic lobe. Development 137, 2981–2987 (2010).

70. A. Androutsellis-Theotokis et al., Notch signalling regulates stem cell numbers in vitro and in vivo. Nature 442, 823–826 (2006).

71. N. Gaiano, J. S. Nye, G. Fishell, Radial glial identity is promoted by Notch1 signaling in the murine forebrain. Neuron 26, 395–404 (2000).

72. K. J. Yoon et al., Mind bomb 1-expressing intermediate progenitors generate notch signaling to maintain radial glial cells. Neuron 58, 519–531 (2008).

73. J. J. Breunig, J. Silbereis, F. M. Vaccarino, N. Sestan, P. Rakic, Notch regulates cell fate and dendrite morphology of newborn neurons in the postnatal dentate gyrus. Proc Natl Acad Sci U S A 104, 20558–20563 (2007).

74. G. Masserdotti et al., Transcriptional Mechanisms of Proneural Factors and REST in Regulating Neuronal Reprogramming of Astrocytes. Cell Stem Cell 17, 74–88 (2015).

75. G. Berdugo-Vega et al., Increasing neurogenesis refines hippocampal activity rejuvenating navigational learning strategies and contextual memory throughout life. Nature communications 11, 135 (2020).

76. N. A. Datson et al., Specific regulatory motifs predict glucocorticoid responsiveness of hippocampal gene expression. Endocrinology 152, 3749–3757 (2011).

77. A. Besnard et al., Targeting Kruppel-like Factor 9 in Excitatory Neurons Protects against Chronic Stress-Induced Impairments in Dendritic Spines and Fear Responses. Cell reports 23, 3183–3196 (2018).

78. J. S. Zhang et al., A conserved alpha-helical motif mediates the interaction of Sp1-like transcriptional repressors with the corepressor mSin3A. Mol Cell Biol 21, 5041–5049 (2001).

79. L. Silberstein et al., Proximity-Based Differential Single-Cell Analysis of the Niche to Identify Stem/Progenitor Cell Regulators. Cell Stem Cell 19, 530–543 (2016).

80. S. L. Otto, J. L. Yakel, The alpha7 nicotinic acetylcholine receptors regulate hippocampal adult-neurogenesis in a sexually dimorphic fashion. Brain Struct Funct 224, 829–846 (2019).

81. A. Carracedo, L. C. Cantley, P. P. Pandolfi, Cancer metabolism: fatty acid oxidation in the limelight. Nat Rev Cancer 13, 227–232 (2013).

82. A. Dobin et al., STAR: ultrafast universal RNA-seq aligner. Bioinformatics 29, 15–21 (2013).

83. S. Anders, P. T. Pyl, W. Huber, HTSeq--a Python framework to work with high-throughput sequencing data. Bioinformatics 31, 166–169 (2015).

84. M. D. Robinson, D. J. McCarthy, G. K. Smyth, edgeR: a Bioconductor package for differential expression analysis of digital gene expression data. Bioinformatics 26, 139–140 (2010).

85. S. Anders et al., Count-based differential expression analysis of RNA sequencing data using R and Bioconductor. Nature protocols 8, 1765–1786 (2013).

